# The Role of Cholesterol in M2 Clustering and Viral Budding Explained

**DOI:** 10.1101/2024.09.09.611993

**Authors:** Dimitrios Kolokouris, Iris E. Kalenderoglou, Anna L. Duncan, Robin A. Corey, Mark S. P. Sansom, Antonios Kolocouris

## Abstract

The proton-conducting domain of the influenza A M2 homotetrameric channel (M2TM-AH; residues 22-62), consisting of four transmembrane (TM; residues 22-46) and four amphipathic helices (AHs; residues 47-62), promotes the release of viral RNA via acidification. Previous studies have also proposed the formation of clusters of M2 channels in the budding neck areas in raft-like domains of the plasma membrane, ^1,2^ which are rich in cholesterol, resulting in cell membrane scission and viral release. Experiments showed that cholesterol has a significant contribution to lipid bilayer undulations in viral buds suggesting a significant role for cholesterol in the budding process. However, a clear explanation of membrane curvature effect based on the distribution of cholesterol around M2TM-AH clusters is lacking. Using coarse-grained molecular dynamics simulations of M2TM-AH in bilayers, we observed that M2 channels form specific clusters with conical shapes, driven by attraction of their amphipathic helices (AHs). We showed that cholesterol stabilized the formation of M2 channel clusters, by filling and bridging the conical gap between M2 channels at specific sites in the N-terminals of adjacent channels or via the C-terminal region of TM and AHs, the latter sites displaying longer interaction time and higher stability. Potential of mean force calculations showed that when cholesterols occupy the identified interfacial binding sites between two M2 channels, the dimer is stabilized by 11 kJ/mol. This translates to the cholesterol-bound dimer being populated by almost two orders of magnitude compared to a dimer lacking cholesterol. We demonstrated that the cholesterol bridged M2 channels can exert lateral force on the surrounding membrane to induce the necessary negative Gaussian curvature profile which permits the spontaneous scission of the catenoid membrane neck and leads to viral buds and scission.

**Significance Statement:** A key role of influenza A M2 channel, a prototype viroporin, is the generation of curvature in the host cell membrane to form budding necks leading to scission and release of the virions. It has been shown experimentally that this process is mediated through the amphipathic helices of M2 and that cholesterol enhanced lipid bilayer undulations in viral budding. Here, using coarse-grained molecular dynamics simulations we revealed that M2 channels form clusters in membranes, induced in an additive manner by both the AHs and cholesterol, which in turn increase local membrane curvature. We showed that both cluster formation and the induced membrane curvature are enhanced by cholesterol by bridging the conical area between M2 channels.

## Introduction

The influenza A M2 proton channel is a canonical viroporin ^3–5^ and is one of 17 proteins encoded by the influenza A viral RNA. ^6,7^ Four M2 peptides (97 amino acids each) assemble with four-fold symmetry into the pH-dependent proton conductance M2 channel, whose (TM) domain forms the pore of the proton channel. ^8–10^ The M2 peptide is a single-pass transmembrane protein with an unstructured N-terminus, a single-pass TM α-helix (M2TM; residues 22-46), ^6,8,11,12^ followed by an amphipathic α-helix (AH; residues 47-62) ^13^ and a highly dynamic C-terminus (residues 62-97). ^14,15^ The M2 channel is blocked by adamantyl amine drugs which bind inside the M2TM pore which has also proton channel properties. ^12,16–20^ However, the extended M2TM-AH, consisting of both the TM and AH domains, forms the conductance domain of the channel (residues 22-62) having specific, amantadine-sensitive proton transport activity indistinguishable from that of full-length M2 (M2FL).^21^ During endocytosis of viral particles, the M2 channel is activated by low pH which causes the imidazole sidechains of the His37 tetrad in the H^+^-pore to be protonated. The electrostatic repulsion between the His37-H^+^ sidechains^22^ drives the M2 channel to adopt an open conformation resulting in proton influx. ^23,24^ The acidification of the viral core leads to the unpacking of the viral genome and to pathogenesis. ^25^ M2TM-AH adopts a wedge- or conical-shaped structure, ^13^ embedding the larger lower half of the cone in the inner leaflet of the biological membrane. This wedge, in turn, generates the required saddle-shaped (negative Gaussian) curvature ^26–33^ required for membrane scission and viral budding. ^34–39^ Thus, the M2TM-AH segment is the functionally important unit of M2, performing both H^+^-channel and membrane-curving roles.

Indeed, recent studies using fluorescence and electron microscopy also found M2 channels to concentrate at the neck of the budding virus in the host’s plasma membrane, ^34–39^ at the boundary between the raft-like and non-raft areas in the membrane, corresponding to L_d_ phases and L_o_ phases. Depending on lipid composition and M2 concentration, the wedge-like M2TM-AH structure allows the channel to both induce and sense curvature, concentrating copies of M2 channels according to biochemical, ^29,32,34,36^ ssNMR, ^27,30,31^ CG MD simulations, ^26,31^ and other biophysical studies. ^27,28,31,33,37,40^ M2TM-AH clustering and membrane curvature effect is stronger in phosphatidylethanolamine (PE) membranes than in phosphatidylcholine (PC) or phosphatidylglycerol (PG) membranes according to ssNMR studies.^33^ Membrane curvature sensing/generation is realized by protein-protein interactions (PPIs) between AHs^41^ as has been reviewed for a plethora of membrane-bound proteins, ^42,43^ including ion channels.^44^ M2 PPIs differ between channel constructs, including bulky hydrophobic TM residues in M2TM clusters, while M2TM-AH clusters did not involve TM domain residues to the same degree, but instead clustered via the AHs as was shown with ssNMR in DOPC/DOPE (DO: 1,2-dioleoyl) lipid bilayers. ^30^ At high local M2 concentrations, as clustering proceeds, the membrane curving effect is exacerbated and influenza A budding begins via the release of the newly constituted virions, ^34–37,39^ attracting more M2 copies to budding necks. Thus, the M2 channel AHs facilitate curvature and budding, ^26–33,45–47^ replacing the need for the endosomal sorting protein complexes required for transport (ESCRT) machinery proteins utilised by other viruses. ^34–37^

Several studies have shown that cholesterol can influence the oligomerization state and energy landscape, e.g., of amyloid-β protein oligomers ^48^ or HIV Fusion Protein gp41. ^49^ Cholesterol significantly contributes to lipid bilayer undulations in viral buds, ^29,50–52^ implying its involvement in the budding process. Similarly, the highly curved neck of an influenza A viral bud is characterised by phase separation involving cholesterol and lipid segregation. ^38,53^ ssNMR ^27,30,31,33^ and CG MD investigations ^26^ have demonstrated that the M2 channel tends to locate itself at the boundary between the raft-like and non-raft areas in the membrane, ^1,2^ where the budding virus can enrich itself with cholesterol to build its viral envelope. ^29,50–52^ While elevated levels of POPE (PO: 1-palmitoyl-2-oleoyl) lipid suppress the capability of M2TM-AH to induce membrane pits, ^51^ cholesterol enhances the ability of M2TM-AH to generate local membrane curvature and pits. ^27,29,31,33,51^ Additionally, atomic force microscopy (AFM) and electron paramagnetic resonance (EPR) spectroscopy, ^51^ fluorescence emission spectroscopy using FITC-M2TM-AH (FITC: fluorescein isothiocyanate), ^54^ Electron Paramagnetic Resonance (EPR) and ^19^F ssNMR ^33^ suggested the influence of cholesterol on the position and orientation of M2TM-AH.

M2-cholesterol interactions are recognised to be specific. Results from ssNMR using labelled cholesterols revealed that cholesterol has a binding site on M2 channel constructs in viral and plasma membrane mimetics including anionic lipids. More specifically cholesterol binding was detected in: (a) M2FL in POPC/POPG/cholesterol bilayers, ^55^ (b) M2TM-AH and M2(22-97) in POPC/POPE/POPS/sphingomyelin (SPH)/cholesterol bilayers (PS: phosphatidylserine). ^56,57^ The cholesterol isooctyl tail interacts with membrane facing Ile39 in M2TM, while the cholesterol polar head lies close to Phe47 at the beginning of AH. Docking predicted that other TM residues such as Leu43 and Leu46 may accommodate the β-face of the hydrophobic steroid core. ^57^ A penta-alanine mutation in AH residues (M2-5Ala: F47A/F48A/I51A/Y52A/F55A) ^29^ led to desensitization of M2 conformational dynamics to cholesterol compared to WT suggesting the importance of the AH domain in cholesterol sensation and conformational regulation. However, the penta-alanine mutation in AH residues was not necessarily budding-defective because of a loss of cholesterol binding. These observations agreed with previous results which showed interaction of cholesterol ^29,51^ deep in the bilayer interface with AHs. Overall, it is now believed that the AH domain interacts directly with cholesterols to facilitate bilayer curvature.

However, it remains unclear how cholesterol is directly involved in clustering of the tetrameric M2 bundles as the mechanism by which the negative curvature neck of the budding virion is formed. CG MD simulations provide a particularly suitable method to explore the interplay between lipid bilayer composition and clustering of M2 channels. Thus, CG MD simulations have been extensively used to investigate the protein-lipid interactions in membranes with cholesterol, ^58,59^ the effect of lipids on the structure and function of membrane peptides and proteins, ^43,58,59^ and the membrane curvature and plasticity, ^41,60–63^ providing significant insights on membrane dynamics and organization that can be linked directly to experiments. ^64–67^ Large-scale CG MD simulations have been also used to investigate protein crowding and its effect on membrane structure and organization. ^43,44^

Previous CG MD simulations showed that M2TM-AH formed linear clusters which generated spontaneous membrane curvature in agreement with experimental findings. ^26–32^ However, M2TM-AH clustering studied by CG MD simulations in refs. ^30^ and ^26^ did not include cholesterol, a major component of the mammalian plasma membrane enriched in raft-like domains and L_o_/L_d_ interfaces. ^1,2,68,69^ Here, we used CG MD simulations to show that M2TM-AH embedded in membranes composed by different lipids, ^51,70^ with 20% cholesterol or without cholesterol present, forms multimeric assemblies. In line with experiment, we observed that in bilayers where cholesterol is present M2 multimers are stabilized that induced membrane curvature. We showed that cholesterol binds between M2TM-AH protomers in specific interprotein clefts situated in the top and bottom membrane leaflets with high occupancies. The top leaflet sites are more dynamic and showed high exchange rates whereas bottom leaflet sites had slower off-rates, favouring the formation of M2TM-AH multimers. We quantified the effect of cholesterol on M2TM-AH clustering via umbrella sampling (US) potential of mean force (PMF) calculations of the dimerization free energies. Cholesterol-induced M2TM-AH clusters increased the Gaussian curvature of the surrounding membrane, a precursor of the budding process.

## Results

### Simulated systems

We applied CG MD simulations for systems including 16 copies of M2TM or M2TM-AH channels in lipid bilayers consisting of 5,000 lipid molecules. In order to explore the effect of M2 clustering ^26,30,33^ and of cholesterol ^28,29,31,33^ on membrane bending we performed CG MD simulations for M2TM channels in DMPC (DM: 1,2-dimyristoyl) and POPC bilayers, the latter without or with 20% cholesterol, for M2TM-AH channels in DMPC, POPC, POPC/POPS bilayers, the latter two without or with 20% cholesterol and for lipid-only bilayers. DMPC and POPC or POPC/POPS ^32,36,37^ bilayers have been applied as membrane models in experimental ^32,36,37^ or simulation studies of M2TM-AH clustering and membrane bending. The POPC/POPS lipids are present in mammalian plasma membranes, which have a negative surface charge due to the presence in the intracellular leaflet of the anionic PS lipids and phosphatidylinositol (PI) lipids e.g., phosphatidylinositol 4,5-bisphosphate (PIP2), ^46^ and have been used as virus mimetic membranes in studies of M2. ^56,57^ We also performed CG MD simulations for M2TM-AH in a plasma-mimetic membrane. The stability of dimers of M2TM-AH channels was tested using atomistic (AA) MD simulations. An overview of our simulations is provided in Supplementary Table 1.

### M2TM and M2TM-AH clustering

In all the membrane systems with 16 copies of M2TM or M2TM-AH tetrameric bundles, we observed that M2 channels clustered, forming dynamic assemblies ranging from dimers to pentamers of the tetrameric channel protein. The scope of this analysis is to predict the kinetics of M2TM-AH cluster formation, starting from the same multiprotein configuration under differing membrane compositions in non-equilibrium conditions. The evolution over time of M2TM-AH clustering in DMPC, POPC, POPC/POPS, POPC/Chol, and POPC/POPS/cholesterol is illustrated in Fig. 1 and for M2TM in DMPC, POPC and POPC/Chol in Supplementary Fig. 1. Simulation repeats (Supplementary Table 1) showed similar results. While M2TM formed a maximum cluster size of four channels in DMPC or POPC or POPC/cholesterol (Supplementary Fig. 1), M2TM-AH formed dimers in DMPC (Fig. 1a) and POPC/POPS (Fig. 1d). In the POPC and POPC/POPS/cholesterol system, M2TM-AH formed dimers to trimers (Fig. 1b and Fig. 1e, respectively) and the biggest cluster – a pentamer – was observed in POPC/cholesterol (Fig. 1c) suggesting cholesterol increases the rate and order of oligomers.

**Figure 1.**
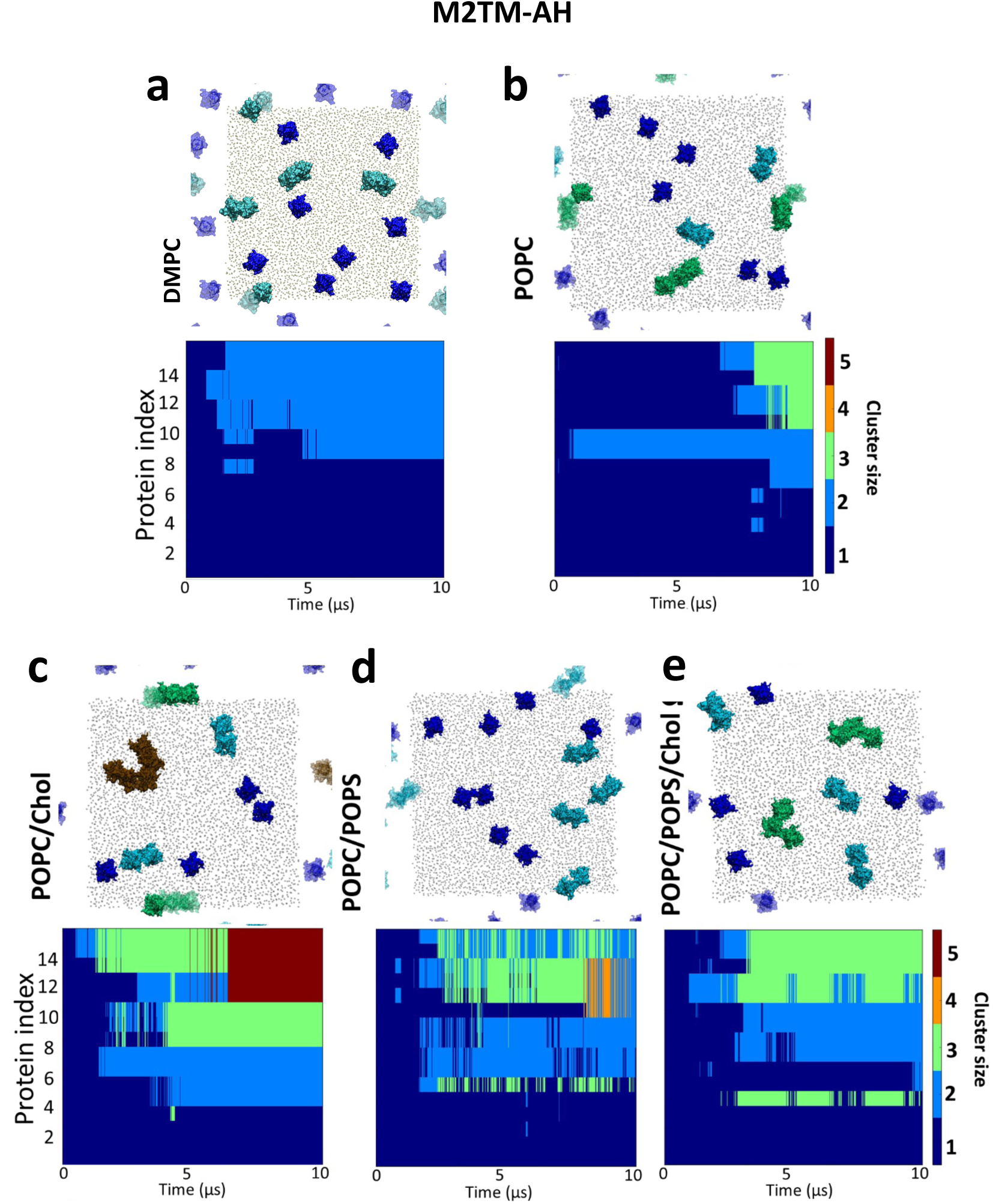
M2TM-AH channel cluster formation over the duration of 10 *μ*s production simulations with the Martini force field ^64,71–73^. The graphs illustrate the time course of M2 channel cluster formation. 16 copies of M2 were simulated in bilayers consisting of **a**. DMPC; **b**. POPC; **c**. POPC/cholesterol (4:1); **d**. POPC/POPS (4:1); **e**. POPC/POPS/cholesterol (3:1:1) (see Supplementary Table 1 for M2TM clustering). The cluster size is indicated by colour (blue for single M2TM-AH protomers, cyan for dimers, green for trimers, orange for tetramers, and brown for pentamers). A representative snapshot at *t* = 10 *μ*s of each system is shown as a top view (*xy* periodicity is shown in transparent).

### M2TM-AH

The shape formation of M2 channel clusters is illustrated in Fig. 2 which shows "slices" of the 16 copy-M2TM-AH channel system from initial (t=0 *μs*; Fig. 2a,c,e,g,i,k,m) and final (t=10 *μs*; Fig. 2b,d,f or o,h,j,l,n) simulation snapshots from CG MD simulations of seven systems in membranes consisting of multimers of M2TM channels in DMPC (Fig. 2a,b) or POPC (Fig. 2c,d) or POPC/cholesterol bilayers (Fig. 2e,f) or multimers of M2TM-AH channels in DMPC (Fig. 2g,h) or POPC (Fig. 2i,j) or POPC/cholesterol bilayers (Fig. 2k,l). The largest M2TM clusters consisting of four M2TM channels were oriented such that their pore axis was approximately parallel to the membrane normal (Fig. 2b, Supplementary Fig. 2). The dimers of M2TM-AH channels were more conical-shaped (see, e.g., Fig. 2h) and thus, the axes of the individual channels were tilted from the membrane normal (Fig. 2h, Supplementary Fig. 2). Interestingly, regarding the pentamer assembly of M2TM-AH in POPC/cholesterol, every M2 protomer was interacting with a maximum of two different M2 channels (linear clustering ^26^). This, in fact, was what we observed in all the clusters of M2TM-AH channels and is likely due to M2TM-AH’s conical shape. Upon the formation of channel dimers, M2TM-AH channel osculated to one another, and their pore axis became tilted to one another. As a result, there was only one binding site for an additional M2 channel. Once a conical shaped dimer of M2TM-AH channels was formed the protein channels generally remained clustered almost throughout the remainder of the simulation.

**Figure 2.**
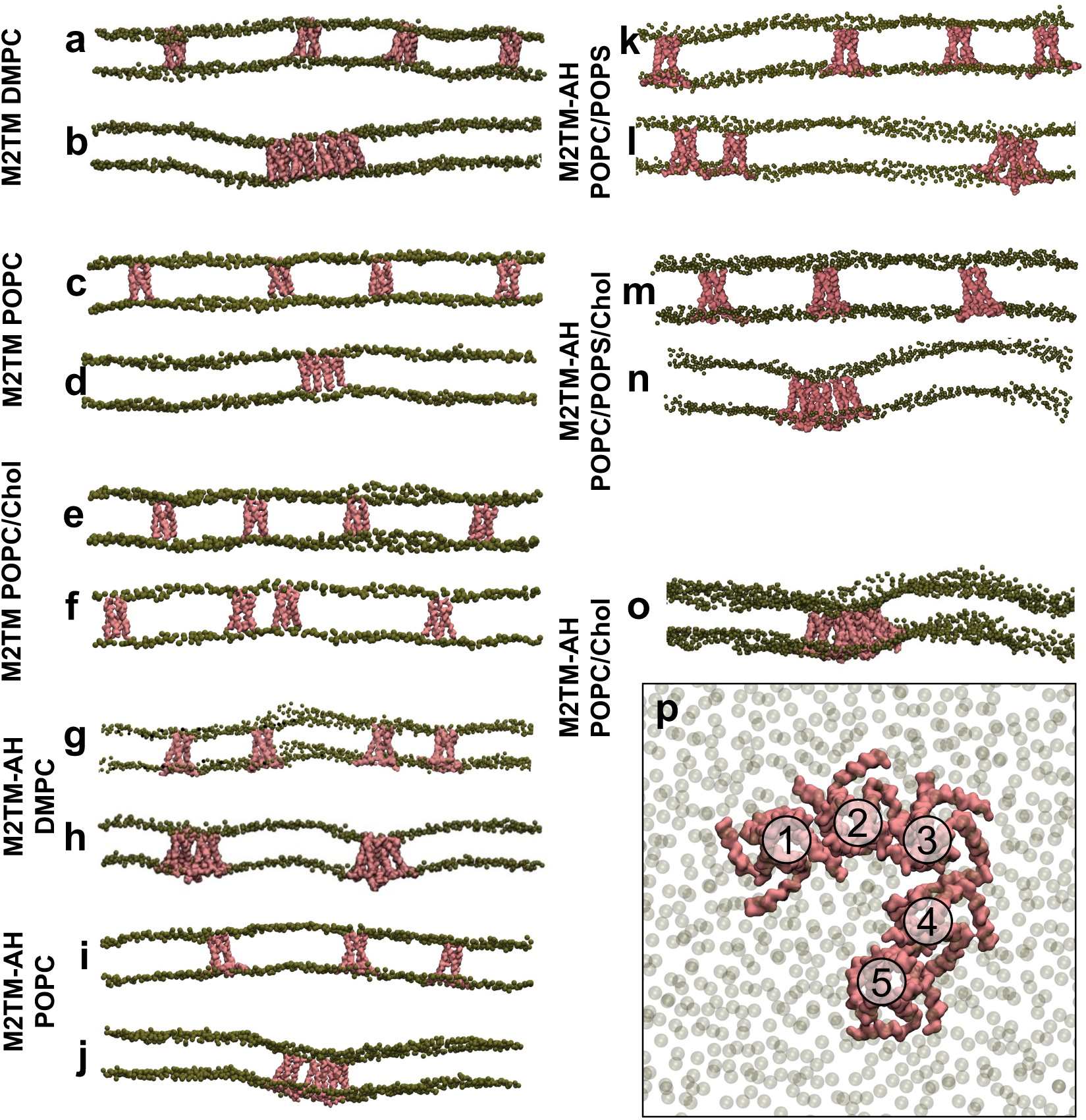
Initial (*t* = 0 *μs*; **a, c, e, g, i, k, m**) and final (*t* = 10 *μs*; **b, d, f, h, j, l, n, o, p**) snapshots of the 16 copies of M2TM or M2TM-AH (backbone beads in pink) in lipid bilayers (phospholipid PO4 beads in green) with Martini force field. ^64,71–73^ The snapshots show **a, b**. M2TM in a DMPC; **c, d**. M2TM in POPC; **e, f**. M2TM in POPC/cholesterol; **g, h**. M2TM-AH in DMPC; **i, j**. M2TM-AH in POPC; **k, l**. M2TM-AH in POPC/POPS, **m, n**. M2TM-AH in POPC/POPS/Chol; **o**. Snapshot at *t* = 10 *μ*s of M2TM-AH in POPC/cholesterol showing the membrane curvature surrounding the pentameric species; **p**. Top view of the M2TM-AH pentamer captured POPC/cholesterol, shown in panel **o**. Each panel demonstrates a clipped view of the full 16-protein copy system. These figures are not representative of the full systems, and they demonstrate only a selected “slice” of the system shown.

### Protein-protein interactions

M2TM-AH channels in the clusters appeared to interact with each other through the residues of the AHs and the N-terminal regions, suggesting that these specific PPIs play a significant role in cluster stabilisation. These findings agree with the previously reported observations in CG MD simulations of M2TM ^30^ and M2TM-AH channels ^26,30^ and with ssNMR results using ^19^F labels. ^33^

We analysed the specific PPIs responsible for clustering of M2 channels by plotting the frequency of the contacts between amino acid residues belonging to different channels. The analysis included all M2 clusters and the replicate simulations. For the M2TM channel clusters the contacts between two adjacent channels included mostly hydrophobic residues throughout the length of the TM domain (Fig. 3a,b). In contrast, for the M2TM-AH channel clusters (Fig. 3c-h) the PPIs between adjacent channels correspond to contacts of polar residues at the N-termini of TM helices and contacts of residues at the C-termini between AHs. ^32–34^ In particular, the residues with the highest interaction frequency between M2TM-AH–M2TM-AH channels in a multimer are the S22 and P25 at the N-termini, and the F54, H57, G58 and R61 of the AHs (Fig. 3c,e,g) compared to the residues located at the transmembrane core (residues 32 to 48). When these PPIs are compared in M2TM-AH multimers in POPC (Fig. 3e,f) and M2TM-AH multimers in POPC/cholesterol systems (Fig. 3g,h), a considerable reduction of the interaction frequencies between TM residues in different protomers is seen in the latter system (Fig. 3e).

**Figure 3.**
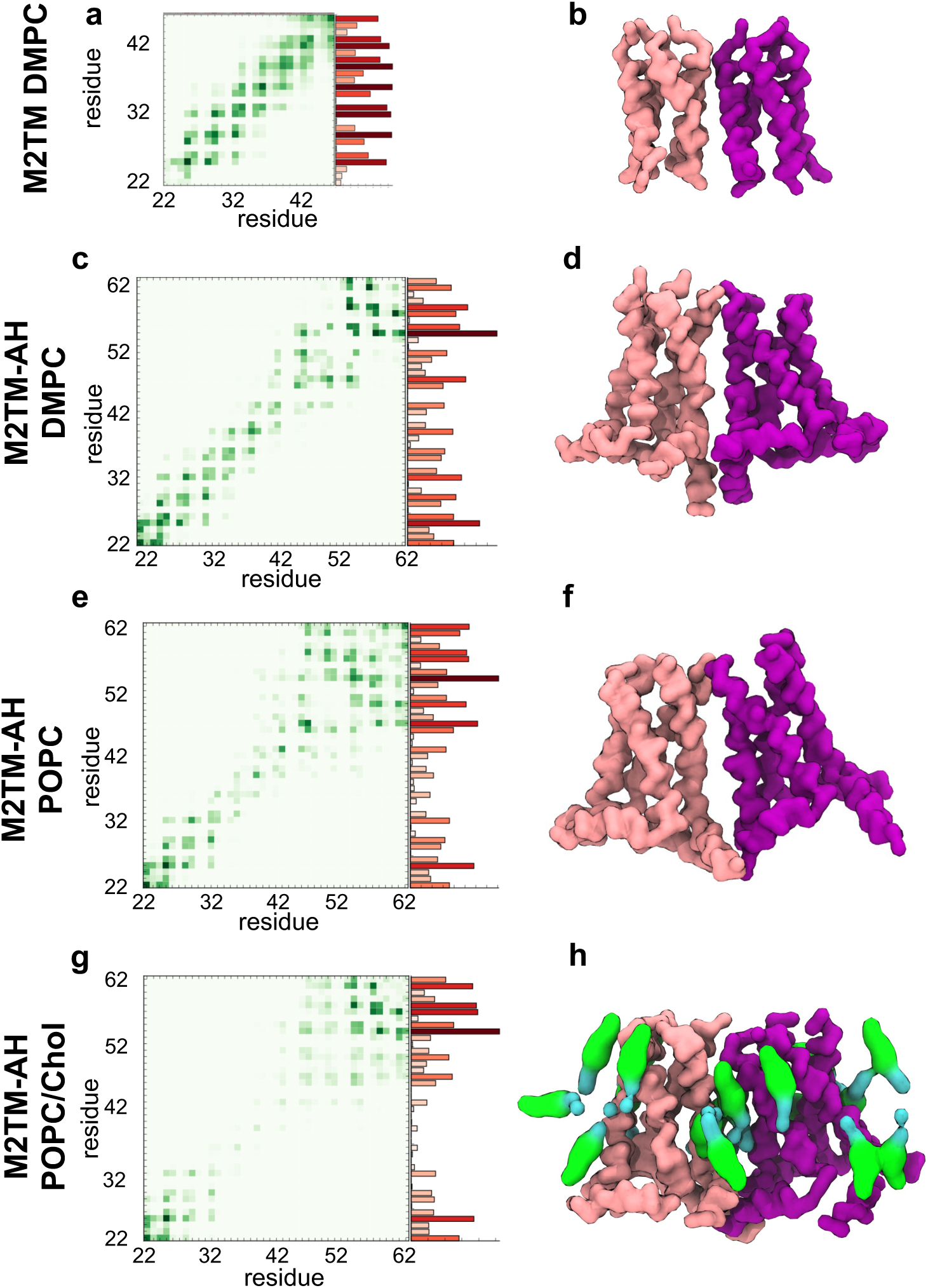
M2TM and M2TM-AH protein-protein interaction (PPI) analysis. The PPIs are presented as interprotomer heatmaps and histograms integrated across the residues of the M2 constructs presented. Data is extracted from 2×10 *μ*s CG MD simulations with Martini force field ^64,71–73^ of 16 M2 construct copies. **a, c, e, g**. Heatmaps correspond to the pairwise interaction frequencies between residues in the adjacent M2 channels shown in **b**, **d**, **f, h,** using a green scale from light green (lower frequency of interactions) to deep green (higher frequency). Histograms show total residue interaction frequency (averaged across all four peptide chains in an M2 channel) using a red scale from light red (lower) to dark red (higher). The representative M2 dimer configurations shown were taken from the last frame of one of the repeat simulations.

Thus, our CG MD simulations suggested that cholesterol molecules are distributed between the TM cores of adjacent channels. This can cause either weakening of multimer formation or cholesterol can act as a bridge between M2TM-AH protomers, potentially acting as an auxiliary mediator. The latter can explain the larger cluster ranks observed in cholesterol-containing membranes (Fig. 1).

### Protein-lipid interactions

We explored the interaction profile of M2 with the bulk lipid environment (Fig. 4; Supplementary Fig. 3). Fig. 4 shows heatmaps of contacts between M2 channel residues with the phospholipid headgroups or cholesterol plotted from CG MD simulations considering all the multimeric states of M2. In the case of M2TM–DMPC or M2TM-AH–DMPC interactions, we analyzed the M2TM– DMPC or the M2TM-AH–DMPC systems, respectively. For the M2TM-AH–POPC, M2TM-AH– POPS or M2TM-AH–cholesterol interactions we analyzed the M2TM-AH–POPC/POPS/cholesterol system. The analysis showed that DMPC or POPC lipid polar heads interacted with the same residues of the M2 protein. Supplementary Fig. 3 shows the M2 amino acid residues that interact with all lipid headgroups; the M2TM and M2TM-AH proteins span the DMPC or POPC bilayer and interacted mainly through the interfacial residues S22, S23, D24 with the polar head region of the top leaflet and mainly through D44, R45, L46 and S50, R53, H57, R61, respectively, with the bottom leaflet (Fig. 4a-c; Fig. S3a-c). ^66^ In the case of POPS, the negatively charged headgroup did not interact extensively with the N-terminal end of M2 which contains the negatively charged D24 residue (Fig. 4d; Supplementary Fig. 3d). For systems of M2TM-AH in POPC/cholesterol or POPC/POPS/cholesterol the interactions between M2TM-AH and bulk cholesterol included the whole protein sequence (Fig. 4e; Supplementary Fig. 3e) with the highest frequency seen for P25. We will analyze in detail the interactions between M2TM-AH and cholesterol in the next paragraphs. The difference in contact frequency between protein and DMPC, POPC or POPS phospholipid head groups (Fig. 4a-d) and the contact frequency between protein and cholesterol (Fig. 4e) was not surprising, because cholesterol extended deep into the membrane and can contact buried residues, several at a time, whereas the phospholipid head groups probably spend most of their time in the interfacial region.

**Figure 4.**
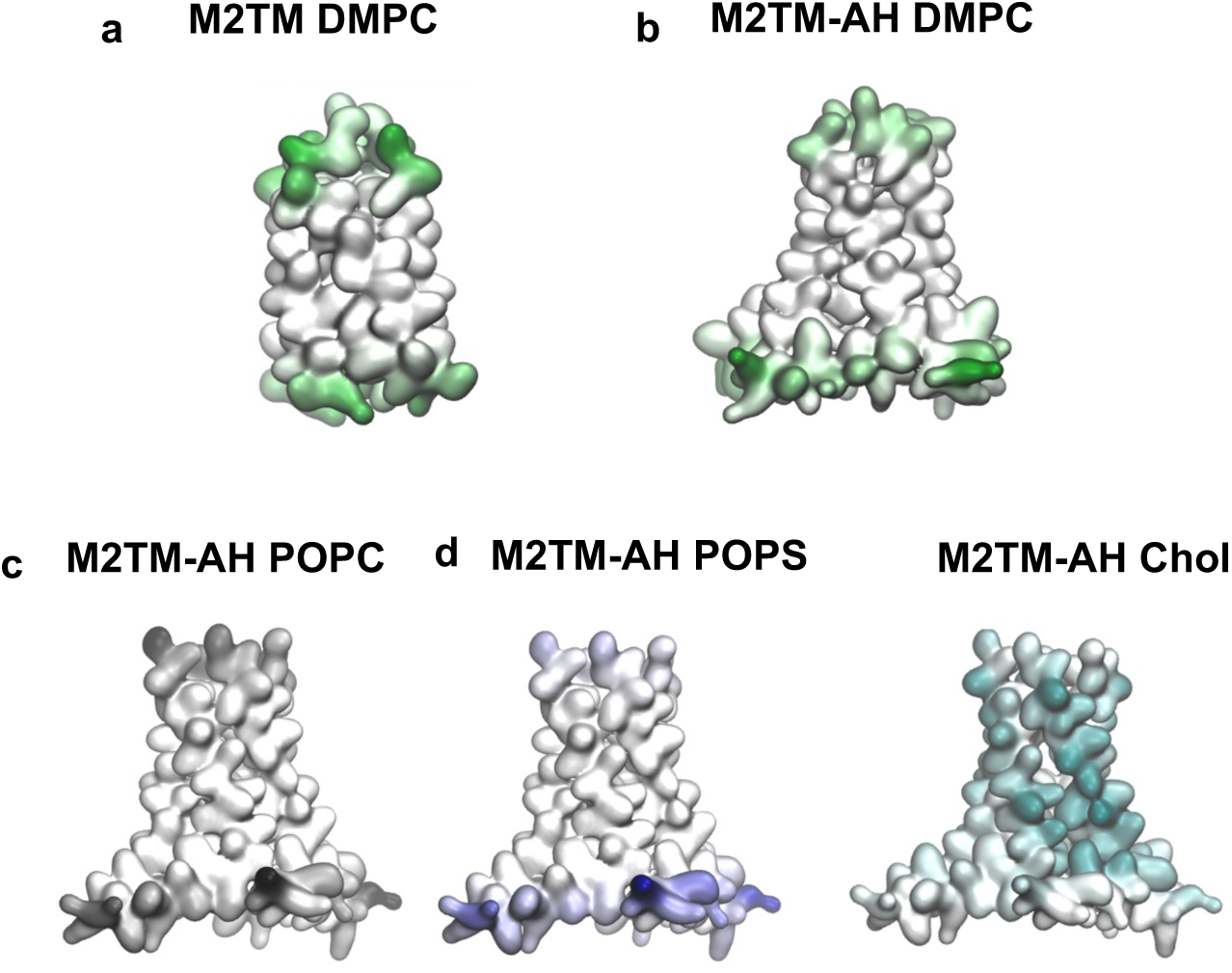
M2TM and M2TM-AH protein-lipid interaction analysis. Lipid interaction heatmaps projected onto CG surface representations of M2TM and M2TM-AH averaged over 2×10 *μ*s and over the 16 independent protein copies per simulated system. Each panel shows data for the indicated lipid type, accounting for interactions with either the phospholipid headgroup (PO4 beads) and/or cholesterol. **a**. M2TM with DMPC lipid headgroups; **b**. M2TM-AH with DMPC lipid headgroups; **c**. M2TM-AH with POPC lipid headgroups; **d**. M2TM-AH with POPS lipid headgroups; **e**. M2TM-AH with cholesterol. For the M2TM–DMPC interactions the M2TM–DMPC system was analyzed. The heatmaps for M2 are coloured according to the interacting lipid; green for DMPC polar heads in **a,b**; grey for POPC polar heads in **c**; blue for POPS polar heads in **d**; teal for cholesterol in **e**.

### Protein-cholesterol interactions

Our CG MD simulations suggested that in POPC/cholesterol and POPC/POPS/cholesterol systems, cholesterol is distributed between the TM cores of adjacent M2TM-AH channels. We observed that the M2TM-AH channel clusters formed include cholesterol molecules between the protomers. To further investigate the equilibrium stability of the clusters formed and their interfacial cholesterols, a patch of membrane with a trimer of M2TM-AH channels was extracted from the POPC/cholesterol system and was subjected to a further 2×50 *μs* MD. This cluster remained a stable trimer during the simulations and cholesterols continued to bridge adjacent M2 protomers (Fig. 5). Calculation of the time-averaged cholesterol density revealed six topologies with pronounced cholesterol density, or, equivalently, three binding sites per degenerate pair of M2 channels (the trimer has C2 symmetry) (Fig. 5a,b). The cholesterol binding sites are shown in Fig. 5a,c. Specifically we report two sites 1, 2 (and their symmetrical equivalents, 4, 3) bridging two protomers situated in the top membrane leaflet. Another cholesterol binding site, 5 (and its symmetrical equivalent, 6) (Fig. 5b,d), in the bottom membrane leaflet.

**Figure 5.**
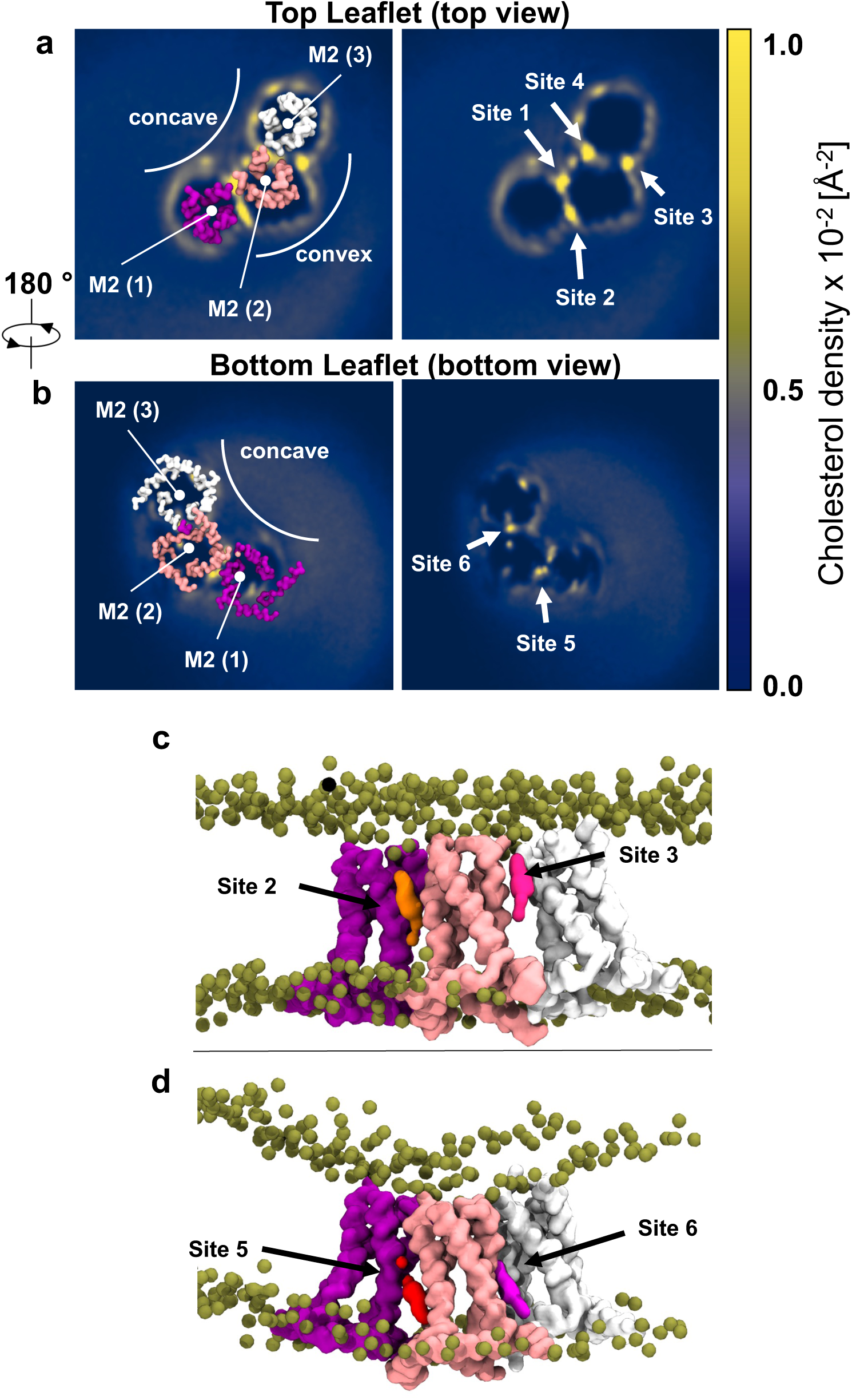
Cholesterol enrichment in the M2TM-AH annulus. 2D time-averaged occupancy of cholesterol shown as membrane-plane projections from 2×50 *μ*s CG MD simulations of a M2TM-AH trimer in POPC/cholesterol (1 Å resolution grid was used). The three M2TM-AH protomers numbered as M2(1), M2(2) and M2(3). **a**. Top view showing the high density – low residency sites 1-4; **b**. Bottom view showing high density – high residency sites 5, 6. Normalized cholesterol density (in Å^-2^) ranges from blue (low values) to yellow (high values); **c**. Side view of a simulation snapshot showing top leaflet cholesterols binding to sites 2 and 3; **d**. Side view of a simulation snapshot showing bottom leaflet cholesterols binding to sites 5 and 6. Proteins are represented as CG surface coloured in purple, salmon, and white (M2(1), M2(2), and M2(3), respectively).

We hypothesized that the aforementioned cholesterol densities may form discrete cholesterol binding sites that stabilize the trimer, and by extension, the M2 clusters which appeared enriched in the presence of cholesterol. For clarity, in Fig. 6 we assign the three M2TM-AH protomers of the trimer as M2(1), M2(2) and M2(3) and present a cartoon schematic of the topology between two M2 protomers M2(1), M2(2) is depicted. Protein-lipid interaction analysis showed that specific residues in the TM α-helices 1-4 (H1-H4) that line the interprotomer space interact with membrane cholesterols (Fig. 6a,b). These residues could be clearly grouped in those interacting with top leaflet and bottom leaflet cholesterols. The four top leaflet cholesterol binding sites of the full trimer (1, 2 or 4, 3) interacted with the N-terminal end of TM in the two channels. Specifically, the per residue interaction analysis showed that in the top leaflet, H1’s high frequency interactions involved residues P25, A29, I32 whereas H4 mostly interacted with cholesterol through residues P25, V28, A29, I32 and I33 (Fig. 6a,b). Supplementary Table 2 shows the extracted kinetic parameters for the three unique cholesterol binding sites for each pair of M2TM-AH protomers calculated using PyLipID. ^74^ Despite the high occupancy of the four top leaflet cholesterol binding sites (sites 1-4; > 65 %), cholesterols showed lower residence time compared to the bottom leaflet sites (sites 5 and 6), suggesting that the bottom leaflet sites have a slower *off* rate compared to those of the top leaflet. Surprisingly, analysis showed different cholesterol binding kinetics between the degenerate pairs top leaflet sites, 1-4 and 2-3. Specifically, sites 1 and 4 have residency 4.7 *μ*s, occupancy 67%, and site surface 7.7 nm^2^ whereas sites 2 and 3 have residency 9.8 *μ*s, occupancy 92%, and site surface 17.1 nm^2^ (Supplementary Table 2). A reduced site surface area is in line with reduced cholesterol accessibility in sites 1-4 compared to 2-3 (Fig. 5a).

**Figure 6.**
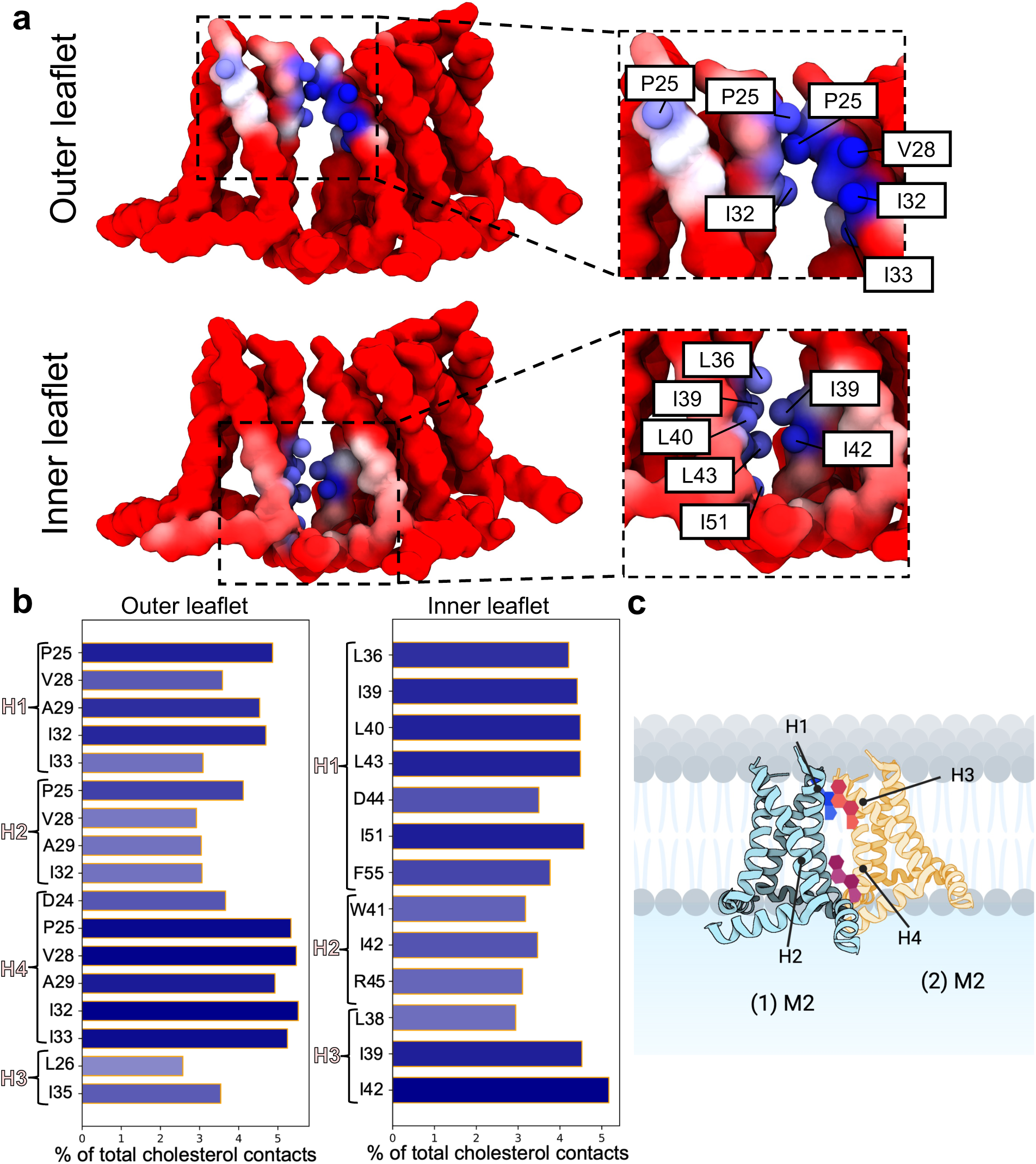
M2TM-AH cholesterol interaction and hotspot analysis **a**. Side view of a snapshot from the 50 *μs* CG MD simulation of the M2TM-AH trimer, with protomers numbered as in Fig. 5. Time-averaged interactions between M2 (1), M2 (2) and top and bottom leaflet cholesterols are analysed. M2(1) and M2(2) residues are coloured according to their interaction frequency with cholesterols, using a red (low) to blue (high) colouring scale. The residues with their side chain beads shown in dark blue correspond to those with the most frequent interactions (> 4% of total cholesterol contacts). **b**. Plots of the most frequent residue interactions from the two M2TM-AH protomers that have contacts with the top and bottom leaflet cholesterols. Contacts are calculated using a 6 Å distance cut-off to define a ‘contact’, based on the radial distribution function for CG Martini ^64,72,73^ lipid-protein interactions. **c**. Schematic representation of the analysis showing two M2 protomers in (a) with their interfacial top and bottom leaflet sites bound by cholesterol. The numbered helices (H) include: H1 and H2 from M2 (1) and H3 and H4 from M2 (2).

The bottom leaflet sites (sites 5 and 6; Fig. 6a) captured cholesterols into specific interprotein binding clefts surrounded primarily by hydrophobic residues with high time-resolved occupancy (88.5 %) and high residence time (15.4 *μ*s) (Supplementary Table 2). This increased cholesterol residency in the bottom leaflet suggested tighter, more specific interactions. At the bottom leaflet residues with high cholesterol interactions frequency are L36, I39, L40, L43, I51 from H1 of the first M2 channel and I39, I42 from H3 of the second M2 channel (Fig. 6a,b). Cholesterol-interacting residues were reported in previous ssNMR experiments ^55–57^ which showed cholesterol binding close to the AH and in contact with residues I39, I42, and F47. We previously captured these residues computationally on a single M2TM-AH showing cholesterol binding (a) in the N-terminal interhelical cleft and (b) between the TM and AH domains (top and bottom leaflets, respectively) with unbiased atomistic simulations and binding kinetics analysis. ^66^ While the lower-residency sites 1 and 4 are on the concave trimer surface, sites 2, 3 and 5, 6 are on the convex surface (Fig. 5a,b) and are associated with cholesterol higher residency times.

To quantitatively weigh the respective contributions of residues coordinating cholesterols at the convex interface we deconstructed the reported binding sites to their residue components and calculated the per residue *off* rates over 2 × 50 μs (Fig. 7). In the top leaflet cholesterol forms hydrogen bonds with S22 (<0.1 *μ*s), D24 (0.13 *μ*s) and are accommodated in the lipophilic cleft by Van der Waals interactions with P25 (4.4 *μ*s), A29 (11.0, 4.2 *μ*s), I32 (8.0, 4.9 *μ*s), V28 (11.4. 2.2 *μ*s). The increased residence time for the latter, hydrophobic residues suggests that the pocket behaves as a “greasy” patch, as suggested by others ^75,76^, without a specific, high-affinity residue. Similarly, in the bottom leaflet, we find cholesterol is engulfed by an extensive lipophilic pocket with the longest, most-stabilizing interactions including TM residues L26 (3.8, 4.7 μs), I35 (5.3, 3.4 μs), L36 (5.3, 5.4 μs), H37 (8.0 μs), I39 (4.2, 3.8 μs), L40 (18.6, 3.1 *μs*), I42 (6.1 μs). In contrast to the top leaflet site, a larger network of hydrophilic interactions coordinate the polar hydroxyl group. Specifically, D44 (1.9 μs) and R45 (1.1 μs) but primarily AH residues S50 (6.3 μs), I51 (2.6 *μs*), F55 (8.2 *μs*) are involved. The residues marked with high binding residency times (Fig. 7) agree with those with high interaction frequencies averaged over all equivalent M2 residues (Fig. 6b). Collectively, the analysis suggests that an extended membrane-facing residue network lines the cholesterol sites. Most importantly, while these residues can make transient interactions with cholesterol in monomeric channels^66^, upon clustering they create “greasy” pockets for steroid group stabilization with more polar residues serving to auxiliary stabilize the hydroxyl group.

**Figure 7.**
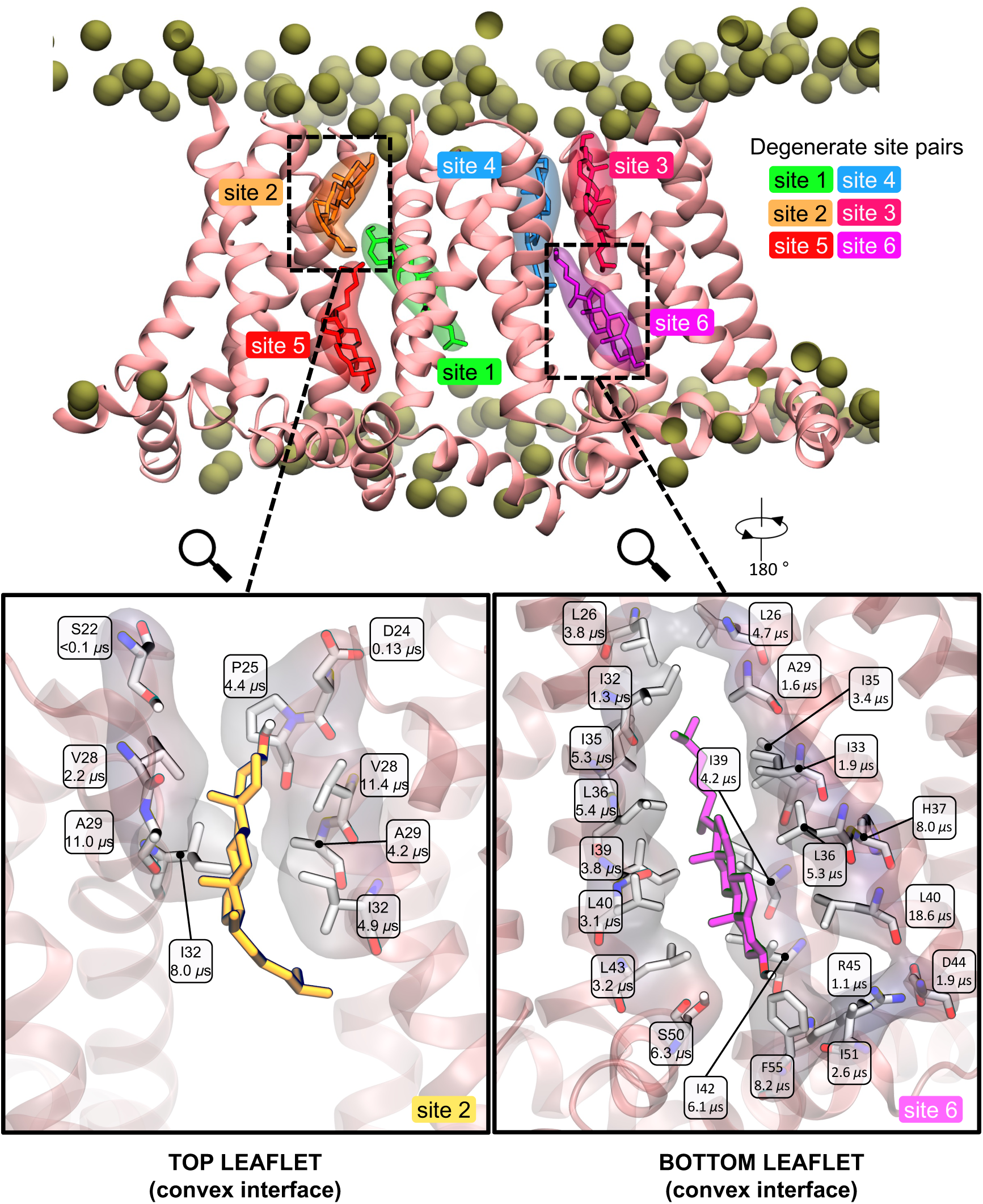
Representative binding configuration of interfacial cholesterols on both leaflets, top (binding sites 1, 2, 3, 4) and bottom (binding sites 5, 6) in 4:1 POPC:cholesterol from 2×50 *μ*s CG MD simulations. The binding configurations were analyzed with the PyLipID python package (see Methods). The binding configuration shown has all high-residency sites occupied (defined as those with overall site residence time > 1.0 *μ*s). Zoomed-in views show the cholesterol-interacting protein residues in white stick representation and transparent off-white Van der Waals surface from binding sites 2 (left) and 6 (right). The per-residue residency time (calculated as 1/*k_off_*, see Methods) is shown. M2 cartoon helices are shown in salmon.

### Energetics of cholesterol-mediated M2TM-AH channel dimerization

Driven by the above results which demonstrated increased multimeric clustering of M2TM-AH in the presence of cholesterol and that cholesterol has well defined interfacial binding sites between clustered M2TM-AH protomers, we proceeded to quantify the relative role of cholesterol in the M2TM-AH dimerization free energy profiles. Dimerization free energies for membrane proteins are computationally demanding and thus scarce in the literature. ^77–81^ We used a M2TM-AH dimer with three cholesterols in between the protomer pairs (in binding sites 1, 4 and 5) from the simulation in POPC/cholesterol bilayers (see Methods). We also considered an equivalent control M2TM-AH dimer lacking cholesterol isolated from the simulations in POPC bilayers. Both dimers (± cholesterols) were embedded and equilibrated in their bilayers and their stability was assessed with the CHARMM36m force field ^82^ to ensure both configurations were stable with an atomistic force field. We verified both dimer configurations to be stable over the course 3 × 500 ns repeats by tracking the interprotomer distance and rotation of one protomer relative to the other over time (Supplementary Fig. 9). Thus, we proceeded with CG MD PMF(US) calculations.

We report both M2TM-AH dimerization free energies (± cholesterols) as PMF(US) profiles calculated using the previously applied weak orientational restraint method to preserve the orientation of a protomer relative to the other ^83^ (see Methods Section) (Fig. 8). Satisfactory US window overlap along the interprotomer distance CV was achieved by inspecting histogram overlap (Fig. S4, S5) and increasing umbrella force constant/reducing window spacing as necessary (see Methods). The dissociation of the dimer of M2TM-AH channels into protomers was driven by linearly increasing the *xy* plane distance between the protomers’ centre-of-mass (COM) from 30.0-56.0 Å. Each US window was equilibrated for 200 ns in NPT ensemble followed by 4.8 *μ*s production time, for a total of 518.4 *μ*s window sampling time.

**Figure 8.**
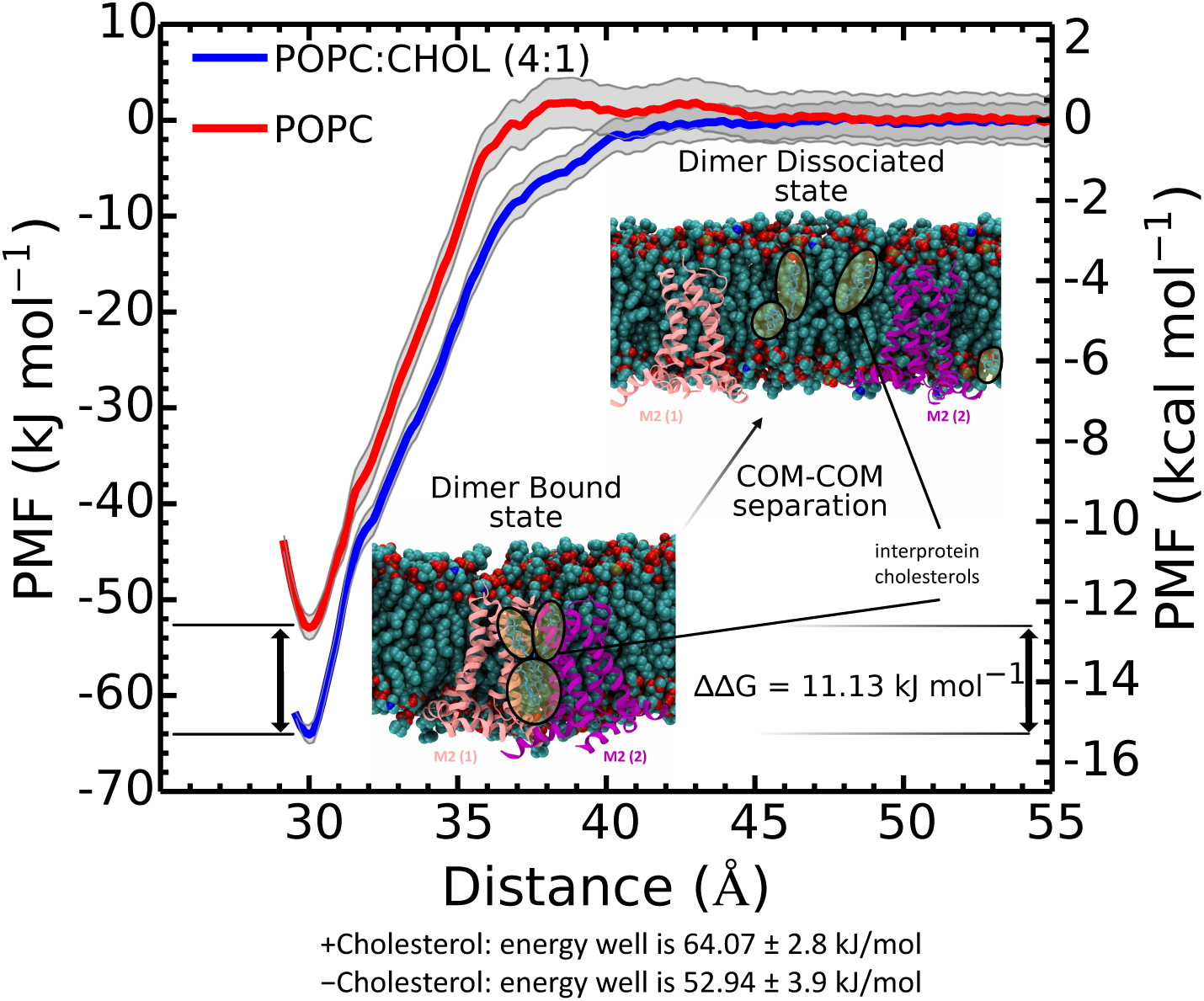
M2TM-AH dimerization PMF profiles (± cholesterols) from CG MD simulations with the Martini force field, ^64,71–73^ analyzed with WHAM. ^84,85^ The PMFs were calculated by dissociating the M2TM-AH protomers from dimers isolated from the larger 16-copy systems in POPC or POPC/cholesterol (4:1). The dimer dissociation into M2TM-AH protomers was driven by linearly increasing the distance between the protomers’ COM on the membrane *xy* plane at a constant rate 0.0001 nm/ns and with a force constant 1000 kJ/mol/nm^2^. The COM distance collective variable ranged from 30.0-56.0 Å and system replicas were selected every 0.5 Å, for a total of 54 windows per system. For the US windows, the force constant was 10 kJ/mol/nm^2^ for 3-4.5 Å and 1000 kJ/mol/nm^2^ for 4.5-5.6 Å. Each US window was equilibrated for 200 ns in NPT followed by 4.8 *μs* production time, for a total of 518.4 *μs* window sampling time (see Supplementary Fig. 4, 5). The calculated free energy well with cholesterol was −64.07 ± 2.76 kJ/mol and without cholesterol −52.94 ± 3.85 kJ/mol.

The PMF landscapes show that within the distance between the COM range of 29-31 Å a local minimum is observed for both systems (Fig. 8). The dimerization free energy of M2TM-AH protomers without cholesterol was –52.94 ± 3.86 kJ/mol (–12.66 ± 0.92 kcal/mol). The dimerization free energy of M2TM-AH protomers with the three interprotein cholesterol sites occupied was –64.07 ± 2.76 kJ/mol (–15.31 ± 0.65 kcal/mol). Thus the PMF(US) calculations revealed that cholesterol strengthens the free energy of association for M2TM-AH assemblies by ΔΔ*G*_POPC→POPC:cholesterol_ = – 11.13 kJ/mol (–2.65 kcal/mol). Since ΔΔ*G* is logarithmically related to the ratio 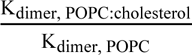, this translates to the M2TM-AH–M2TM-AH dimer state being almost 2 orders of magnitude more populated when the interprotein sites are occupied by cholesterol. This result agrees with our observation from unbiased MD of increased M2 cluster sizes upon addition of 20% mol cholesterol in the membrane (Fig. 1b,d). Cholesterol binds between two adjacent M2TM-AH protomers in a dimer in sites with high occupancy and slow *off* rates. Collectively, this suggests that cholesterol acts as a molecular glue by strengthening M2TM-AH association via specific protein-cholesterol-protein bridges, thus enhancing multimer formation.

We also note that the PMFs are in agreement with the out-of-equilibrium time-dependent oligomer tracking presented in Fig 1 for POPC vs POPC/cholesterol. Specifically, the increased dimerization energy of M2TM-AH when the interfacial pockets are occupied with cholesterol (Fig 8) qualitatively agrees with the higher oligomerization speed observed when cholesterol is present in the membrane (Fig 1b and c).

### M2AH-cholesterol complexes influence membrane curvature

The AHs of M2 channel mediate the membrane undulations ^26–33,51^ which are necessary to the membrane-curvature mechanism of virus budding. ^34–38 26–33,51^ While multiple single point mutations in the AHs did not affect viral replication suggesting that specific residues are not important ^86^, the penta-Ala M2 construct, where five bulky hydrophobic residues on the AH are replaced with alanines, is known to promote significantly reduced negative Gaussian curvature^28^. Furthermore, previous studies suggest that cholesterol enhances the ability of M2TM-AH to generate membrane curvature. ^26–33,51^ We show above that clustering of M2TM-AH enhances the deformation of planar membranes during unbiased MD simulations (Fig. 2). By extension, we turned our focus on quantifying the curvature produced by M2TM-AH in our model membranes and whether this is affected by the absence of the AHs or cholesterol. An initial analysis showed that time-averaged local curvature cancelled out over time (partly due to the diffusive behaviour of proteins in bilayers) meaning that we were unable to use average membrane curvature across the simulation time as a metric. However, visual inspection of the trajectories suggested differential membrane curving between M2TM-AH, M2TM and protein-free membranes (Fig S10 a-j). Membrane deformation along the *z*-axis (membrane normal) showed that M2TM-AH clusters are populated in membrane valleys (as seen from a top view, Fig S10 b,f,g,h), whereas M2TM is able to cluster outside valleys as well (Fig S10 a,c,d). We note that while a protein-free POPC bilayer remained relatively planar (Fig S10 i), the addition of cholesterol caused spontaneous deformations (Fig S10 j), as also seen in a recent study where cholesterol was found to be enriched in areas of increased curvature^87^.

We hypothesised that clusters of M2TM-AH channels could exert a linactant deformation force to annular lipids and via surface tension modulation regulate curvature. To quantify M2’s effect on membrane planarity, we calculated the Gaussian curvature (*K*) on a surface fitted to the membrane, z*_ij_* (*i* and *j* define grid cells (*i,j*)), defined as 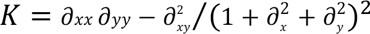, and projected on the membrane *xy* plane (Fig 9a-i, Fig S6). The data were averaged over the final 2 *μ*s of the simulations as longer averaging times result in “flattening” of the landscapes due to undulations traversing the *xy* plane. *K*>0 means a spherical membrane protrusion (on either side of the membrane; i.e. hills or valleys) whereas *K*>0 refers to a saddle-shaped hyperboloid. We also tracked the time evolution of the absolute net Gaussian curvature (|*K*|) over the whole simulation (10 μs) as a measure of global membrane curving (Supplementary Fig. 7). Equivalent *K* heatmaps resulting from the second simulation repeats are shown in Fig. S11. Both repeats were averaged and shown in scatter dot plots (Supplementary Fig. 6) and statistics are tabulated in Supplementary Table 4. Additionally, *K* heatmaps averaged over only the final 100 ns of the repeats accentuate the temporally resolved membrane undulations (Supplementary Fig. 12, 13).

**Figure 9.**
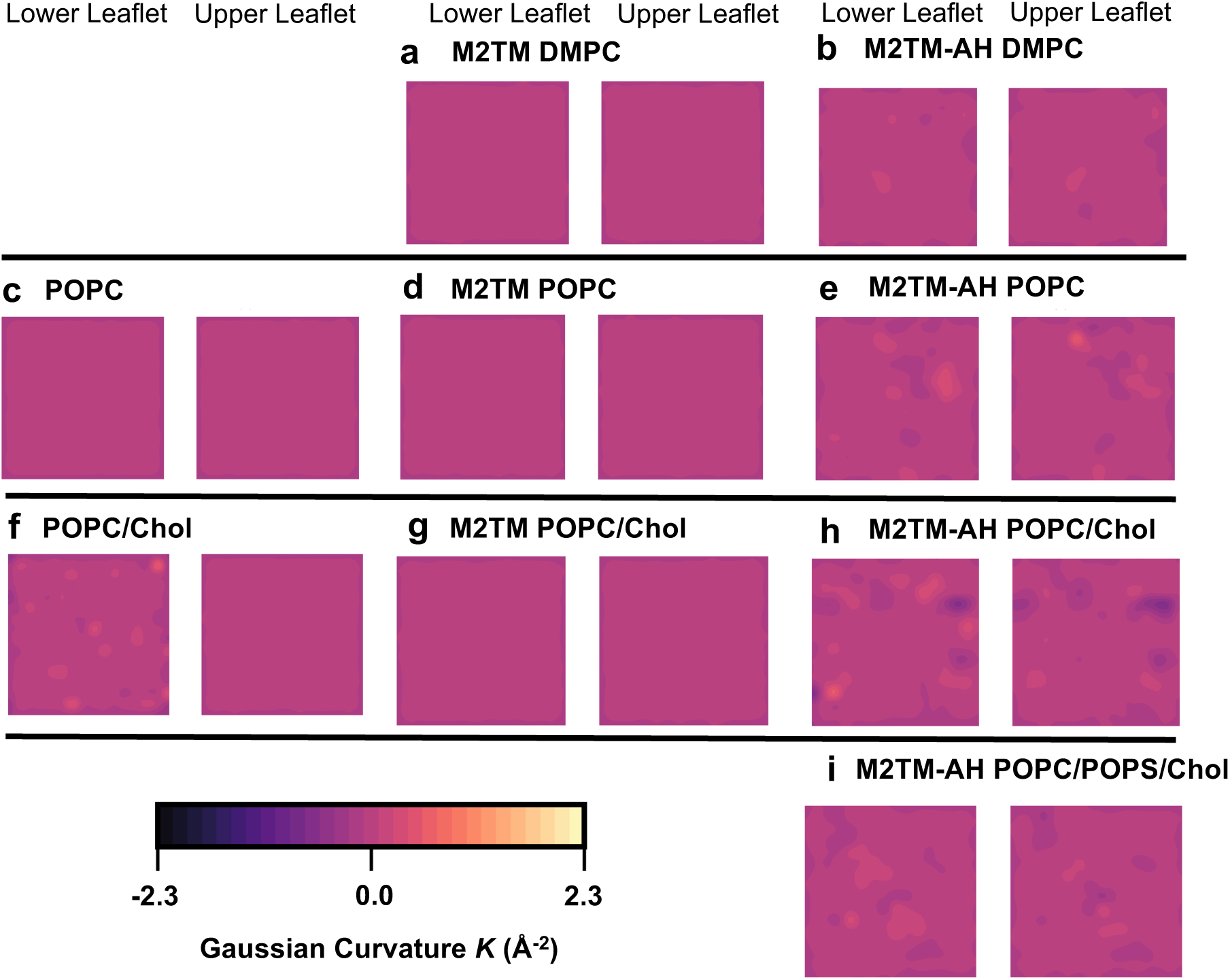
Gaussian curvature (*K*) induced by M2 constructs (±AHs) and cholesterol from 10 *μ*s CG MD of 16 copies of the M2 constructs and protein-free control bilayers with the Martini force field. ^64,71–73^ The *K* heatmaps are averaged over the final 2 *μ*s from a single replicate. Simulated systems shown are **a**. M2TM in DMPC; **b**. M2TM-AH in DMPC; **c**. POPC; **d**. M2TM in POPC; **e**. M2TM-AH in POPC; **f**. POPC/cholesterol; **g**. M2TM in POPC/cholesterol; **h**. M2TM-AH in POPC/cholesterol; **i**. M2TM-AH in POPC/POPS/cholesterol. *K* is shown in a colour gradient ranging from black to bright yellow (−2.3 to 2.3 Å^-2^). Results for the second replicate are shown in Supplementary Fig. 11. Averaging over shorter simulation intervals highlighted the quicker, temporally resolved *K* fluctuations. Examples over the last 100 ns are shown in Supplementary Fig. 12, 13.

The analysis showed that when AHs are present, local membrane curvatures are clearly enhanced compared to the curving observed in M2TM systems lacking the AHs (Fig. 9). The effect of M2TM-AH increased the Gaussian curvature range of a POPC bilayer (protein-free; −0.221 to 0.035 Å^-2^, +M2TM; −0.253 to 0.034, +M2TM-AH; −0.302 to 0.577 Å^-2^) (Supplementary Table 4). Interestingly, the AHs did not however exert such a pronounced effect on a stiffer DMPC membrane (+M2TM; - 0.272 to 0.296, +M2TM-AH; −0.241 to 0.211 Å^-2^) suggesting that M2TM-AH’s curvature induction is limited by the bending modulus (25.3 and 34.7 k_B_T for POPC and DMPC, respectively^88^). We also find that cholesterol increased the global curvature in both leaflets when added in a protein-free bilayers (protein-free POPC; −0.221 to 0.035 Å^-2^, protein-free POPC/cholesterol; −0.570 to 0.589 Å^-2^), in agreement with ssNMR results ^89^ and CG MD simulations reported by others. ^87^. Adding M2TM-AH in POPC/cholesterol induced further negative and positive *K* in both leaflets (Fig. 9h), being the largest *K* range between all the systems compared (+M2TM-AH; −1.357 to 0.952 Å^-2^). Notably the AHs coupled with cholesterol produced the most negative *K*, the curvature type topologically necessary for a number of membrane destabilization events including membrane budding and scission. Removing the AHs only modestly increased system-wide curving (M2TM/POPC/cholesterol; −0.245 to 0.046 Å^-2^, M2TM/POPC; −0.253 to 0.034 Å^-2^), suggesting that cholesterol’s curvature inducing effects are mitigated in the presence of just the TM domain. Our result highlights that when both components are present spontaneous saddle-shaped patches can be formed without external bending forces.

The quantitative comparisons presented here suggest an additive, synergistic mechanism between the AHs and cholesterol, consistent with their roles in increased clustering. (Fig. 1c). Clustering in POPC/cholesterol membranes is localized in curved membrane patches, where *K* is non-zero (Fig. 2o and Fig. S10f). This system showed the largest M2 channel cluster sizes (reaching up to pentamers) and the most stable in time clusters (Fig. 1c), with trimers forming within the first *μ*s of the simulation repeats. From a membrane biology standpoint this suggests that the AHs are essential to navigate M2 channel clusters to curved membrane patches where cholesterol is enriched^87^ or, vice versa, that the AHs are essential for the clusters to induce local curvature.

Addition of the anionic POPS in the M2TM-AH/POPC/cholesterol mixture suppressed negative Gaussian curvature (Fig 9j, +POPS; −0.322 to 1.046 Å^-2^, −POPS; −1.357 to 0.952 Å^-2^). Despite the reduction in *K* range, M2TM-AH forms higher-order oligomers in the presence of POPS (>2 channels; Fig. 1e). To better understand this, we tested whether the interfacial cholesterol binding sites we discussed above (see protein-cholesterol interactions section) are still able to bind in the presence of POPS. Indeed, when we embedded and simulated the isolated trimer in a membrane that includes both cholesterol and POPS, the interfacial cholesterol sites were still identified (Supplementary Table 3). The analysis showed residency times >1.5 μs and occupancies >50% with the addition of POPS, showing that interfacial cholesterol can bind the same sites. However, we noted decreases binding residency times for the interfacial sites (see Supplementary Table 2 versus Supplementary Table 3). We wondered whether this might be due to a direct POPS competition effect for the interfacial sites. However, no interprotein POPS binding sites were found. A single non-interfacial, peripheral POPS binding site was found (not shown) with lower residency time (0.84 μs) and low occupancy (32.6%) over three simulation repeats, suggesting that the reduction in cholesterol binding observed is non-competitive. Other POPS hotspots showed <0.1 μs binding times and <25% occupancies suggesting that compared to cholesterol, POPS is dynamically mobile with distinct POPS molecules exchanging rapidly. Thus, POPS reduces *K* and exerts a non-competitive reduction in interfacial cholesterol binding (in the concentration used, 20% mol, i.e. 1:1 mol with cholesterol). Notably, POPC/POPS without cholesterol also resulted in transient M2TM-AH cluster formation/deformation over time (Fig. 1d), which we did not observe in the other lipid systems. ^87,89^

## Discussion

For many enveloped viruses, including HIV and influenza, assembly and budding occur from membrane microdomains (e.g., lipid rafts) ^36,90^ simultaneously with virion formation. Understanding the mechanisms underlying this process could promote biomedical efforts to block viral propagation.

M2 preferentially localizes at the edge of membrane rafts ^1,34–38,44^ which develops into the bud neck position (also referred to as the budozone). ^34–39^ Such a localization of a fission-inducing protein at the neck of the budding virus near lipid rafts ^1,2^, causes membrane undulations in model planar membranes. ^26–33,51^ Previous studies have suggested the formation of clusters of influenza A M2 channels in the saddle-shaped budding neck resulting in cell membrane scission, suggesting possible involvement of M2 clustering in the viral budding mechanism. ^26–38^ The AHs of M2 were shown to contribute to cluster formation and membrane undulations in planar plasma membranes, ^38^ facilitating viral budding. ^34–39,41^ However the clustering mechanism remained unresolved. The M2 AHs utilize the steep dielectric at the lipid-water interface and the water concentration gradients (between the hydrophobic core and lipid headgroup region) to produce membrane undulations ^26–34,51,70^ and increase lateral pressure during budding. ^35,91^ Both M2TM or M2TM-AH insert deeply into the bilayer and induce the formation of an ordered lipid domain. ^35,36,38,51,92^ Increased lipid ordering and generation of the suitable membrane curvature at the neck adds further strain on the constricted, phase-separated membrane neck, enabling additional constriction and causing membrane scission during *in vivo* budding. ^34–39,41^ However, the catenoid-shaped (*K*<0) influenza budding neck has a diameter of 25.97 ± 11.25 nm, ^37^ i.e. much larger than that of spontaneous necks arising due to M2 in SUVs (4.67 nm). ^28,70^ [Note: this calculation assumed similar bending rigidity between the SUVs and the membrane neck of a budding virus possibly underestimating M2’s constricting capacity *in vivo*]. Nevertheless, it has been suggested that the scaffolding-induced constriction and phase segregation alone are insufficient to cause membrane scission in the absence of M2. ^32,34,37,38,41^ Rather, the combination of scaffolding-induced constriction and curvature induced by M2 is sufficient to cause membrane scission. ^28,37,38,70^ Therefore M2 may be sufficient to cause scission in pre-constricted necks, ^32^ by reducing the neck diameter by an additional 5 nm via exerting Gaussian curvature of −0.04 nm^−2 28^ at the midpoint of the neck catenoid. ^32,93^

^26–35,51,70,86,35,86^In addition to the native membrane asymmetry, the distribution of cholesterol in the budozone raft-like domain ^1,2,53,68,69^ influences protein organization. There have been several observations that cholesterol impacts M2-driven membrane remodelling. There is significant association between elevated membrane curvature and reduced amantadine binding with the AH-cholesterol interaction. ^31,94^ Eliminating either component (AH or cholesterol) diminishes membrane curvature, thereby affecting the packing of the transmembrane helix and reducing drug accessibility to the pore. ^31,94^ Increased bilayer thickness, a known cholesterol effect, also leads to a more compact homotetramer. ^95^ As a result of cholesterol’s condensing properties, ^96^ the amphipathic helix shifts away from the membrane in response to cholesterol supplementation. However, cholesterol is not essential for the association of the amphipathic helix with membranes (LUVs), ^97^ nor for M2 ion channel function and inhibition by rimantadine ^98^ challenging the assumption of its functional necessity. Another suggested mechanism leading to protein clustering involves cholesterol inducing a shift in the conformational equilibrium of M2 toward a predominantly α-helical state. ^99,100^ This implies a potential function in promoting interactions with the helical M1 protein. ^38^ However, in low-cholesterol membranes, the tilted M2TM-AH peptide increases lipid order and likely increases membrane line tension. ^54^ High cholesterol levels enhanced separation of anionic lipid headgroups, and the membrane-parallel insertion of M2TM-AH protein channel reduces headgroup separation without affecting lipid order. ^54^ Additionally, M2, influenced by cholesterol, induces negative membrane curvature, especially in virus budding lipid raft sites. ^37^

It has been shown that cholesterol interacts with M2TM-AH. ^29,55–57,66^ ssNMR with labelled cholesterol revealed proximity to residues I39, I42, and F47 of the M2TM-AH channel and a stoichiometry 2:1 for the cholesterol–M2TM-AH complex. Moreover, cholesterol contribution in M2TM-AH clustering has been suggested to be mediated via direct M2TM-AH–cholesterol contacts based on distances between M2TM-AH channels using ^19^F ssNMR. ^33^ More macroscopically, membrane protein crowding has been known to induce large spontaneous curvature.^101^ However, there was no direct interpretation at a molecular level of the cholesterol mediated M2 clustering and membrane undulations.

In the present work we aimed to (a) provide a microscopical description as to how cholesterol is mediating M2 channel clustering with subsequent effects of clustering on membrane morphology, and (b) to provide molecular insight to previous experimental findings. MD simulations can contribute to the understanding of viral structure, functional dynamics, and processes related to the viral life cycle.^102^ Here, we provided a model of multimeric formation of M2TM-AH and the membrane curvature induced by M2TM-AH in planar membranes. Using 10 *μs* repeats of CG MD simulations of M2TM-AH and M2TM in DMPC, POPC, and POPC/POPS without or with cholesterol, we showed that M2TM-AH channels are clustered ^26,30,33,37^ forming multimers through PPIs between AHs in adjacent channels. M2TM that lacks the AHs formed clusters through direct attraction between parallel M2TM helices. The stabilizing PPIs between M2TM-AH protomers are formed by contacts between polar residues at the N-termini of the TM helices and contacts between residues at the C-termini of the AHs in adjacent channels (Fig. 3). The multimers that are formed during the simulation are mainly dimers, trimers (and even pentamers) while the supplementation of cholesterol in as a membrane component favours the formation of higher order multimers (Fig. 1, 2). In POPC/cholesterol and POPC/POPS/cholesterol all the clusters have cholesterol molecules bridging paired M2TM-AH channels (Fig. 3, 5, 6, 7). Indeed, our PMF(US) calculations of M2TM-AH channel dimerization in POPC and POPC/cholesterol membranes (Fig. 8) showed ΔΔ*G*_POPC→POPC:cholesterol_ = 11.13 kJ/mol (−2.65 kcal/mol) which corresponds to M2TM-AH-M2TM-AH being populated by a factor of 71 compared to monomer species. Cholesterol acting as a molecular glue makes the binding between two M2 proteins tighter and multimeric clustering is increased.

In terms of lipid composition, we used symmetric planar bilayers composed of DMPC or POPC as reference membrane models, in order to investigate the specific effects of cholesterol or anionic lipids (POPS). In the field of M2, and in general when membrane proteins are reconstituted in proteoliposomes, the experimental data typically use simple lipid compositions due to the practical difficulties in asymmetric self-assembly of bilayers. Examples include ssNMR structural studies of M2TM in DMPC^18^ (where DMPC matches with M2TM’s hydrophobic length), or M2TM-AH in lipids with longer aliphatic chains (e.g. DOPC/DOPE) ^13^ or M2FL in DMPC/DMPG bilayers. ^55^ The membrane remodeling effect by M2TM-AH was studied using ssNMR and CG MD simulations in DOPC/DOPE bilayers ^30^ and the M2TM-AH channels clustering affected by cholesterol was studied using ssNMR in POPC/POPG/cholesterol or POPE bilayers.^33^ In other studies the bending effect in membranes by M2TM-AH and M2TM was investigated in DOPS/DOPE or DOPS/DOPE liposomes using small-angle X-ray scattering (SAXS), ^28^ or the M2TM-AH in POPC:cholesterol or POPC/POPG/cholesterol liposomes using an array of biophysical techniques. ^32^ The experimental work done so far with model membranes aims to describe cholesterol effects in viral budding. However, the basis has not been described using models that enable molecular-level interpretation. Taking this data into account, we performed our computational experiments using symmetrical lipid mixtures composed of selected key lipid types known to influence curvature and viral budding. Our system selection lies in between models including natural asymmetric lipid membranes and a reductionist *in vitro* biochemical approach, with the aim to interrogate the role of cholesterol.

Nevertheless, alongside our symmetric bilayer compositions (reflective of those used in experimental setups), we have also used a more native-like asymmetric plasma membrane model (see Methods Section; includes different leaflet PC/PE distribution, relatively high cholesterol and higher levels of glycosphingolipids) and used this to expand our analysis. We postulated that the lipid makeup could influence protein clustering dynamics. We found that M2 molecules were able to cluster in a plasma membrane mimetic bilayer creating higher order (>2) oligomers compared to the simple, symmetric POPC environment clustering is limited to dimers in our analysis of 10 *μ*s repeat simulations (see Supplementary Fig. 8). Notably, monosialodihexosylganglioside lipids (GM3) tend to create a tight perimeter around M2 copies, increasing interprotomer distance. The presence of the lipid annulus complicates the analysis of protein-protein interactions preventing us from analyzing them in the same way as for the simpler membrane models, requiring different PPI interaction cutoffs to be optimized, and ±GM3 simulations. However, the stability of protein complexes remains evident, as illustrated by the consistent distance between the COMs of M2TM-AH channels (Supplementary Fig. 8c). The clusters in the plasma membrane mimetic bilayer are similar to those formed in the symmetric POPC/cholesterol and POPC/POPS/cholesterol environments (Fig. 1). Whilst studies on GM3 as an annular lipid are mostly restricted to MD studies, it has been shown experimentally that GM3 does form clusters in the membrane. ^103,104^ It becomes evident that protein-lipid and potentially lipid-lipid interactions can affect clustering outcomes, especially when considering that ganglioside GM3 lipids display increased affinity for M2 by forming a tight lipid annulus. Physiological GM3 could therefore further augment M2 clustering by dragging M2s in GM3-enriched membrane patches. This phenomenon could be explored in more depth, necessitating an analytical approach that takes the lipid annulus into consideration. As this would require extensive work beyond the scope of the present analysis we reserve this for a future manuscript, especially since GM3’s effect on M2 membrane dynamics is entirely unexplored. Concluding, it is important to recognize the role of lipid composition when exploring protein-lipid-protein interactions. A path for future M2 clustering experiments could be geared towards emulating membrane complexity, by using both lipid extracts and chemically defined complex bilayers.

We calculated the time-averaged density of bridged cholesterol molecules from CG MD 2×50 *μs* of a trimer of M2TM-AH channels in POPC/cholesterol membrane and we identified three cholesterol binding sites bridging M2TM-AH channel pairs with distinct residue interactions (Fig. 5, 6). Definition of binding sites with a recently published community analysis method identified two sites of high occupancy located towards the M2TM-AH N-termini in the top leaflet, ^66^ not experimentally observed previously. ^55–57^ Importantly the two top leaflet sites differed in their membrane accessibility, with the site exposed to the convex trimer interface geometry (Fig. 5) showing higher occupancy and cholesterol residency compared to the one in the concave interface. Another cholesterol site of high occupancy/high residency (i.e., tight binding and slow exchange with bulk) was predicted at the C-ends and AHs in the bottom leaflet that has been observed by ssNMR. ^55–57^ The binding site of cholesterol in the bottom leaflet is more stable compared to the binding sites in the top leaflet and the analysis of the per residue residence time showed that in the bottom leaflet cholesterol forms polar interactions with D44, R45, S50 and have very frequent hydrophobic contacts with L26, I35, L36, H37, I39, L40, I42, I51, F55. Conversely, the top leaflet binding sites are stabilized through supporting polar interactions bonding with S22, D24 and frequent van der Waals interactions with P25, A29, I32, V28. The specific contribution of each site residue was ranked with per-residue residence time (Fig. 7).

We report that M2 clustering and membrane curvature is favoured by cholesterol in agreement with previous findings. ^33–37,50–52,105^ Furthermore, PPIs between adjacent M2 channels can be divided into two categories: (A) PPIs involving interacting amino acid residues located at the N- and C-termini, and (B) cholesterol-mediated PPIs, referred to as cholesterol bridges, which enable protomers to come into close proximity and form larger clusters. The latter mechanism involves a reduction in the free energy of dimerization mediated by cholesterol bridges. Specific cholesterol binding sites attract cholesterol molecules to fill the gap between the hydrophobic cores of two adjacent M2s and promoting M2 dimerization.

Upon examining the effect of lipid-tail length on M2TM and M2TM-AH channels clustering, it became apparent that M2TM channels demonstrated a higher propensity for clustering in comparison with M2TM-AH channels in DMPC (Fig. 1, Supplementary Fig. 1). On the other hand, when considering POPC, M2TM-AH channels exhibited higher clustering levels, while clustering of M2TM channels is significantly diminished. This observation suggests the existence of a lipid-sensitive packing effect on M2TM, particularly when AHs are present. As discussed in our previous work, in which we investigated M2TM and M2TM-AH protomers in different lipid environments through AA MD simulations, ^66^ both systems exhibited membrane thickness convergence to the same value regardless of the construct. Therefore, M2’s adaptation mechanism to adjust to different membrane thicknesses (i.e. M2TM in POPC or M2TM-AH in DMPC) might involve structural changes that hinder either protein diffusion or aggregation. Consequently, it is of paramount importance to carefully select the combination of M2 construct and lipids when studying M2’s clustering mechanism. Since the choice of the M2 construct plays crucial a role in the clustering mechanism, as shown here and by others ^26,30,33,37^, further investigation is needed including M2FL in various membrane compositions.

We observed that the AHs induce greater local deformation compared with M2TM channels in planar lipid bilayers (Fig. 2, Supplementary Fig. 10). The clusters in M2TM-AH systems assist valley formation or are trafficked to existing membrane valleys spontaneously or both in a synergistic manner. Cholesterol enhances the formation of multimers and valleys. In contrast, in the equivalent M2TM simulations, clusters are located both in valleys and hills of the membrane and there is no obvious correlation between cluster *xy* position and local membrane curvature (Supplementary Fig. 10). We measured the Gaussian curvature (*K*) that M2 channel constructs can generate on the membrane (Fig. 9) and found that while the lack of AHs (M2TM) resulted in time-averaged planar membranes, the addition of AHs on M2 is enough to generate multiple negative Gaussian curvature events with *K*< –0.04 nm^-2^ across the membrane plane. The effect is pronounced with the addition of cholesterol, demonstrating that the steroid can link multiple aspects of M2 physiology by (a) binding to selected high and low exchange rate sites between M2 protomers to create (b) protein-cholesterol-protein bridges that thermodynamically favor dimer populations by 1.5-2 orders of magnitude, which in turn (c) drives channel clustering into conical multimer shapes that exert lateral force on the membrane to induce (d) the necessary negative Gaussian curvature profile to permit spontaneous scission of the catenoid membrane neck and lead to (e) viral buds and scission.

It is difficult to discern from our analysis whether it is the membrane curvature that primarily drives M2 channel clustering or whether M2 channels clustering in the presence of cholesterol bridges can create conical protein formations which force the membrane in a curved shape. Thus, in other studies using CG MD simulations and generic lipid bilayer models ^106,107^ it was suggested that once a minimal local bending by a protein is realized, the effect robustly drives protein cluster formation and subsequent transformation into vesicles with radii that correlate with the local curvature imprint. ^108^ In these simulations, when small proteins were placed in a simulated lipid environment, similar in scale to lipid microdomains, vesicular structures formed spontaneously despite a lack of inter-protein interactions. Rather, the vesicularization was induced by local hydrophilic attraction between the proteins and their immediate lipid environment. These local attractions led to aggregation and local membrane curvature which summed over the membrane to induce vesicle formation. For example, it was found that matrix (M) protein of the vesicular stomatitis virus alone was able to impose the correct budding curvature on the membrane using confocal microscopy in giant unilamellar vesicles (GUVs) consisting of pure DOPC or from a mixture of DOPC/DOPS (9:1). ^109^ Additionally, when virus capsid-sized particles were placed in the lipid environment, these also induced spontaneous vesicle formation.

Notably by the time this manuscript was in the revision process, cholesterol parameters for Martini 3 were published. ^110^ The utilization of the MARTINI force field in simulation studies has yielded valuable insights into the association of membrane proteins ^77,79,80,111–114^ and protein-lipid interactions. ^59,115,116^ It has been reported that the MARTINI 2.2 model used has certain limitations, ^117,118^ notably in the smoothing of free energy landscapes, which may have led to accelerated oligomerization kinetics. MARTINI 2.2 utilizes a shifted function with a cutoff at 1.1 nm and implicit screening to depict electrostatic interactions, instead of relying on the particle mesh Ewald approximation. This decision may lead to the potential underestimation of long-range electrostatic interactions between proteins. Consequently, interactions involving charged lipids and the protein through electrostatic forces could be overestimated. Nevertheless, the M2 channels clustering was also previously observed in CG MD simulations ^26^ using the BPB force field. ^119,120^ Additionally, it is important to emphasize that the key lipid of interest in this study is neutral cholesterol, and its interactions occur within relatively short distances (less than 1.1 nm). A just published work reported the reliability of cholesterol Martini 2.2 parameters for cholesterol based on the agreement between simulations and experimental results. ^87^ Importantly, our CG MD simulations results agree with the experimental evidence suggesting cholesterol molecules in between M2TM-AH channels. ^33^ Nevertheless, subsequent investigations into M2 oligomerization using MARTINI 3 ^121^ are poised to be intriguing. Additionally, after the submission of this manuscript, Kim et al. demonstrated that for very large membrane systems (100 nm × 100 nm × 80 nm, >30,000 lipids and >6.5 million waters) the default neighbour list update step and interaction cutoff, *r*_c_, used in MARTINI 2.2 simulations need to be refined to avoid artificial curving of a lipid membrane.^122^ Our simulated systems are smaller (40 nm × 40 nm × 10 nm, <5,000 lipids, <80,000 waters) and we did not observe excessive artificial curving in protein-free control POPC bilayers (Fig 9 and S6).

## Conclusions

Cholesterol is the molecular ingredient commonly associated with protein lateral diffusion and mobility. ^2,123,124^ It is possible that the lateral diffusion rates of M2TM-AH protomers and associated viroporins through the membrane to engage other protomers/dimers/trimers etc. are affected in the presence of cholesterol thereby allowing rapid translocation of the channels across the *xy* plane. Using CG MD simulations, we found that M2 channels clustering and membranes curvature is favoured by cholesterol in agreement with previous findings. ^33–37,50–52,105^ The higher order multimers promote membrane curvature in the cholesterol dense membrane during viral budding. Cholesterol concentration can play a dual role in M2 mesoscale dynamics by balancing (a) the viroporin’s dimerization affinity and (b) the probability of multimerization. It is likely the latter proceeds via a membrane fluidity modulation mechanism, which cholesterol is known for, but this remains to be tested. The simulations presented here inform our understanding of the mesoscale organization of influenza A M2 channels in membranes of the host cell during virus budding. These findings may have implications for the design of novel medications for influenza virus infection via inhibition of cluster formation and will trigger related research for other viruses. ^49^ It has been suggested that membrane remodelling can contribute to virus budding effected by the matrix protein 1 (M1) of influenza C,^125^ the non-structural protein 1, and the Ebola Virus Matrix Protein VP40. 126 The SARS-CoV-2 homo-pentameric protein E forms clusters 127 and also causes membrane bending. 128,129 It has been suggested for HIV that cholesterol increases clustering of viroporin gp41 channels wherein cholesterol bridges multiple gp41 trimers, increasing proximity and facilitating virus-cell fusion and virus budding. 130,131

## Methods

### System setup for CG MD simulations

We performed the simulations shown in Supplementary Table 1. For the M2TM CG MD simulations we used as starting structure the final snapshot obtained from a 200 ns-AA MD simulation starting from the experimental structure of M2TM with PDB ID 2KQT. 18 We converted the AA model of M2TM to a CG model using the *martinize.py* script (version 2.4), available on the MARTINI coarse grain force field website (http://md.chem.rug.nl/index.php/tools2/proteins-and-bilayers). We then applied the MARTINI2.2 force field 64,72,73,132 and the ElNeDyn elastic network model. 112 The CG model of M2TM channel was embedded into a membrane bilayer of the desired lipid composition which was solvated using the standard MARTINI water model 64,72,73,132 and neutralized to the physiological 0.15 M NaCl concentration using the *insane.py* script 133 available on the MARTINI website. This system of M2TM channel in a simulation box dimension with 10 nm × 10 nm × 10 nm, was minimized, equilibrated and simulated for 1 *μ*s. We used the final snapshot of the M2TM channel system to assemble a larger system of 4 × 4 copies of the M2TM channel, using the GROMACS 134,135 tool genconf, consisting of 16 proteins and ∼ 5000 lipids. This system of 16 M2TM channels was subjected to CG MD simulations for 10 *μ*s.

For the 10 *μ*s CG MD simulations of M2TM-AH systems, we also started from the final snapshot obtained from a 200 ns MD simulation of M2TM-AH embedded in a hydrated lipid bilayer consisting by DMPC or POPC or POPC/POPS, ±20% cholesterol, starting from the experimental structure of M2TM-AH with PDB ID 2L0J. 13

We performed GG MD simulations testing M2TM-AH channel clustering in a plasma membrane mimetic model. The extracellular leaflet of the plasma mimetic membrane is enriched in phosphatidylcholine (PC) lipids while the intracellular leaflet is enriched in phosphatidylethanolamine (PE) lipids and phosphatidylserine (PS) lipids. Thus, the upper leaflet consisted of 4:4:1:1:3:2:5 POPC: DOPC : DOPE : POPE : SPH : GM3 : cholesterol while the lower leaflet was 1:1:4:4:2:1:2:5 POPC : DOPC : POPE : DOPE : POPS : DOPS : PIP2 : cholesterol.

To perform the 50 *μ*s CG MD simulation starting from a trimer of M2TM-AH channels in POPC/Cholesterol, we used a trimer of M2TM-AH channels with cholesterol molecules bridging the protomers from the last snapshot of the 10 *μ*s CG MD simulation of M2TM-AH in POPC/cholesterol; in this trimer each cholesterol molecule was between the protomers in a distance of 9 Å between the COM of cholesterol and the COM of each M2 protein. We extracted this trimer from the above-mentioned trajectory using VMD. 136 The trimer of M2TM-AH channels was embedded into a POPC/cholesterol 20% membrane bilayer, solvated using the standard MARTINI water model 64,72,73,132 and neutralized to a 0.15 M NaCl concentration (Supplementary Table 1), using the *insane.py* script. The resulted system had a simulation box dimensions 12 nm × 14 nm × 10 nm, and was minimized, equilibrated, and subjected to CG MD simulations for 50 *μ*s repeats.

### CG MD simulation parameters

We performed all simulations using GROMACS 5.1.2 (www.gromacs.org) 134,135 and the MARTINI 2.2 force field. 64,72,73,132 In all CG MD simulations, we applied periodic boundary conditions and a time step of 20 fs. The temperature was maintained at 320 K using the Berendsen thermostat 137 and the pressure was maintained at 1 bar using the Berendsen barostat, 137 except from the simulation of the trimer of M2TM-AH channels, where a velocity rescale thermostat 71 and Parinello-Rahman barostat 138 were used. A coupling constant of 1 ps was used for temperature coupling and a coupling constant of 10 ps was used for pressure coupling. The electrostatic interactions were calculated using a coulomb-type potential and for van der Waals interactions a switching function from 0.0 to 1.1 nm was used and a cutoff distance at 1.1 nm. LINCS algorithm was used. 139 All systems were subjected to CG MD simulations for 10 *μ*s repeats except the trimer of M2TM-AH channels system which was simulated for 50 *μ*s repeats.

### Backmapping and AA MD simulation parameters

We carried out backmapping of the CG MD simulation snapshots to AA models in CHARMM36m force field 82 using CG2AT2140.

For selected systems (see Supplementary Table 1), AA MD simulations were performed using the CHARMM36m. 82 Initial structures were obtained by transferring the equilibrated structures from the CG MD simulations to atomistic resolution with approximately 168,000 atoms. The temperature was kept at 320 K via the v-rescale algorithm. 71 Isotropic pressure coupling to 1 bar was applied using the Parrinello-Rahman algorithm 141,142 with a compressibility of 4.5 *·* 10^−5^ bar^−1^, Van der Waals forces were smoothly switched to zero (between 1.0 nm and 1.2 nm), and the electrostatic interactions were treated with the PME method. 143 The time step was 2 fs. The all-atom systems were studied at a salt concentration of 0.150 M NaCl. The systems were subjected to AA MD simulations for 500 ns repeats.

### Cholesterol binding site identification and analysis

The 50 *μ*s trajectory of the isolated trimer of M2TM-AH channels in POPC/cholesterol system was analysed with the PyLipID 74 python library. In PyLipID 74 the cholesterol-interaction cut-offs specified were 0.475-0.75 nm. Cholesterol occupancy and density visualisation was carried out with VMD 1.9.4. 136 We selected a representative trimer frame that captured the average cholesterol profile (*t* = 8.116 *μ*s) for further analysis. This was done by binning the calculated cholesterol occupancy to an isosurface of isovalue 0.08 and superimposing the 50 *μs* trajectory (VMD 1.9.4 136 VolMap tool), frame-by-frame.

### Protein-Protein, Protein-Lipid Interactions and Clustering Analyses

PPIs and protein-lipid interactions were identified using in-house clustering scripts, making use of the NumPy 144 and MDAnalysis 132 modules. Residues of neighbouring proteins were considered as interacting when residue centroids were within 0.8 nm of one another. Similarly, for the clustering analysis, in-house scripts using the NumPy, MDAnalysis and NetworkX Python libraries were used. In the clustering scripts, M2 channels were considered to interact with each other when the centres of mass of the whole protein were within 3.9 nm of each other (for M2TM-AH) and 2.7 nm (for M2TM). Graphs were plotted using xmgrace, gnuplot 4.6 (www.gnuplot.info), Matplotlib 145 and for molecular visualization VMD 136 was used. For protein-lipid contact analysis, a cut-off distance of 0.6 nm was employed, based on radial distribution functions for CG lipid molecules. 146 The 2D density map was computed by using the VolMap VMD plugin tool. The occupancy density calculations were performed by applying a grid over the simulation box with 1 Å resolution and were averaged across all frames and normalised. Rotation and translation motion of the protein was alleviated by fitting the protein backbone using trjconv -fit option in GROMACS as well as the VMD RMSD alignment tool. Scripts used to analyse PPIs can be found at: https://github.com/annaduncan/clustering_prot and scripts to analysis protein-lipid interactions at: https://github.com/annaduncan/Kir_scripts/blob/master/lipid_prot_interaction_frequencies_v5.py

### Umbrella sampling PMF calculations

We performed all protein-protein PMF calculations 83 with GROMACS 134,135 2020.3. The systems consisting of dimers of M2TM-AH channels were prepared for PMF(US) calculations as follows. A representative frame from the 50 *μ*s CG MD simulations trajectory (*t* = 8.116 *μs*) of the trimer of M2TM-AH channels was selected with all interfacial cholesterol sites occupied. The trimer of M2TM-AH channels consists of two adjacent topological dimers. One of the two topological dimers in POPC/cholesterol was selected for subsequent PMF calculations, by removing the coordinates for the third M2TM-AH protomer. After the removal of a M2TM-AH protomer, the truncated system was supplemented with ions to reach 150 mM NaCl and charge neutrality. The M2TM-AH_dimer_ system was subjected to two rounds of steepest descents minimization followed by bilayer self-assembly after 1 *μ*s equilibration to fill the membrane gap caused by the removed protomer. The self-assembly step was done in NPT with a conservative 10 fs timestep and with remaining parameters as in the CG MD simulations section above. The equilibrated dimer of M2TM-AH channels in POPC/cholesterol system was used to prepare an identical system without cholesterol. 113 cholesterol coordinates were deleted and the dimer of M2TM-AH channels in POPC system was equilibrated by another round of bilayer self-assembly. Position restraints were applied on the protein backbone in the dimer of M2TM-AH channels using MARTINI beads (*k_x-y plane_* = 1000 kJ/mol/nm^2^), thus yielding starting channels dimer coordinates identical to the +cholesterol system.

Having carefully prepared comparable dimer of M2TM-AH channels in POPC/cholesterol and POPC bilayers, with or without cholesterol in between the M2TM-AH protomers, the starting configurations were used in steered MD to drive dissociation. The dissociation of the dimer of M2TM-AH channels into protomers was driven by linearly increasing the *x-y* plane distance between the protomers’ COM at a constant rate 0.0001 nm per ps and with a force constant 1000 kJ/mol/nm^2^. The COM distance collective variable ranged from 30.0-56.0 Å and system snapshots were selected every 0.5 Å, for a total of 95 windows per system. The PMF curves plateaued to bulk values within the selected range. For the US windows, the force constant was 10 kJ/mol/nm^2^ for 3-4.5 Å and 1000 kJ/mol/nm^2^ for 4.5-5.6 Å. Orientational restraints were applied to the protomers to prohibit undesired rotation about the M2TM-AH *z* axis which would increase time to convergence. This was achieved by applying a rotational potential to the particles of each protein. This is equivalent to setting the rotation matrix Ω(*t*) elements to 1 for particle entries. The potential chosen does not influence the positional umbrella potential, since it has been proved 147 that radial forces and forces parallel to the rotation axis are eliminated by using the rotational potential form according to:

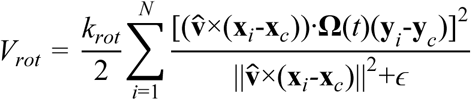

where 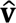 is a unit vector parallel to the rotation axis; **x***_i_* and **y***_i_* are the current and reference positions of the *i*^th^ MARTINI particle, respectively; **x***_c_* and y*_c_* are the current and reference positions of the centre of mass of the *N* MARTINI particles in each protein, respectively; **Ω**(*t*) is a rotation matrix, which describes the motion of the potential; *k_rot_* is the force constant for the rotational potential, and *ε* is a small constant required to avoid a singularity at the axis of rotation. 83

This method has been previously used to calculate NanC dimerization free energies. 83 The rotational potential is applied via the enforced rotation code in GROMACS (rot_type0=rm2-pf in mdp file), which applies an appropriate translation force to each restrained particle to avoid rotational torques. *k_rot_* = 1000 kJ/mol/nm^2^ and *ε* = 0.01 nm^-2^ were selected as previously reported to be appropriate for a membrane protein system. 83 Each US window was equilibrated for 200 ns in NPT followed by 4.8 *μ*s production time, for a total of 518.4 *μs* window sampling time. Within this time the PMFs obtained had converged. To reconstruct the window-mean forces into a PMF curve, gmx wham 84,85 was used with 200 bootstrap rounds, 200 bins and a tolerance of 1e-06. The average profile and error bars extracted from gmx wham were plotted in Python 3 with the matplotlib library.

### Gaussian curvature (*K*) measurements

Gaussian membrane curvature (*K*) was calculated by using in-house python scripts, making use of the NumPy, 144 SciPy 148 and the MDAnalysis 132 module as implemented in their MembraneCurvature package. 149 The phospholipid phosphate beads (PO4) were selected as reference points for grid generation and the average Gaussian curvature was calculated over two independent repeat simulations and over (a) the last 100 ns to better capture the fluctuation range or (b) the last 2 μs to better curvature statistics.

## Data and Software Availability

Information about simulation systems setup, software and scripts used are provided in the Methods section. The simulation input coordinates (gro, pdb files) and parameter files (mdp, itp files) of the CG MD simulations described in Supplementary Table 1 are available via: https://github.com/dimkol94/M2-cholesterol-clustering

## Acknowledgements

DK is supported by the BBSRC studentship number (BB/M011224/1) and the Onassis Foundation PhD scholarship award (F ZO 035-1/2018-2019). IK thanks Erasmus+ program for the traineeship funding her stay at the University of Oxford (7198\2018). IK is supported by the European Union’s Horizon 2020 research and innovation programme (Marie Skłodowska-Curie grant agreement No: 860954). AD is supported by the BBSRC grant number (BB/R00126X/1) and Pembroke College, Oxford (BTP Fellowship). MSPS: BBSRC (BB/R00126X/1), EPSRC (EP/R004722/1; EP/V010948/1), Wellcome (208361/Z/17/Z). AK thanks Chiesi Hellas (SARG No 10354).

## Abbreviations

AH: amphipathic helix
ER: endoplasmic reticulum
ESCRT: endosomal sorting complexes required for transport
CG MD: coarse-grained molecular dynamics
DOPC: 1,2-dioleoyl-*sn*-glycero-3-phosphocholine
DOPE: 1,2-dioleoyl-*sn*-glycero-3-phosphoethanolamine
HCV: hepatitis C virus
HIV: human immunodeficiency virus
M2 protein: matrix-2 protein
MD: molecular dynamics
M2TM: transmembrane domain of M2 protein
POPC: 1-palmitoyl-2-oleoyl-*sn*-glycero-3-phosphocholine
POPE: 1,2-dipalmitoyl-*sn*-glycero-3-phosphoethanolamine
POPS: 1-palmitoyl-2-oleoyl-*sn*-glycero-3-phosphoserine
PMF: potential of mean force
PPI: protein-protein interaction
PV: poliovirus
SARS CoV: severe acute respiratory syndrome coronavirus
US: umbrella sampling
vpu: viral protein U
WHAM: Weighted Histogram Analysis Method

## Author Information

### Author Contributions

AK conceived and designed the research; AK & MSPS supervised the research; IK performed the unbiased CG MD simulations supervised by ALD; DK performed PMF(US) and binding site calculations; IK and DK analyzed the trajectory data and prepared the figures; IK, DK, ALD, MSPS, RAK, AK interpreted the data; IK, DK, AK wrote the manuscript. All authors edited the manuscript.

*IK current address:* Laboratory for Membrane Protein Dynamics, Department of Neuroscience, Faculty of Health and Medical Sciences, University of Copenhagen, Copenhagen, Denmark.

*ALD current address: Department of Chemistry,* University of Aarhus, Aarhus C, Denmark

