## Supplementary Information for "The Role of Cholesterol in M2 Clustering and Viral Budding Explained"

### Supplementary Tables

**Supplementary Table 1.** CG MD-based simulations with Martini force field [64,127,128,135] performed to investigate M2 channels clustering and membrane bending in different lipid bilayers. [a]

| Simulation CG | PDB ID | Number of proteins | Number of lipids | Bilayer composition | Repeats × duration |
|---|---|---|---|---|---|
| MD in POPC (lipid-only) | - | - | 5024 | 1 | 2 × 10 $\mu s$ |
| MD in POPC:cholesterol (lipid-only) | - | - | 5024 | 4:1 | 2 × 10 $\mu s$ |
| MD in M2TM DMPC | 2KQT | 16 | 5024 | 1 | 2 × 10 $\mu s$ |
| MD in M2TM POPC | 2KQT | 16 | 5024 | 1 | 2 × 10 $\mu s$ |
| MD in M2TM POPC:cholesterol | 2KQT | 16 | 5024 | 4:1 | 2 × 10 $\mu s$ |
| MD in M2TM-AH DMPC | 2L0J | 16 | 4800 | 1 | 2 × 10 $\mu s$ |
| MD in M2TM-AH POPC | 2L0J | 16 | 4800 | 1 | 2 × 10 $\mu s$ |
| MD in M2TM-AH POPC:cholesterol | 2L0J | 16 | 4768 | 4:1 | 2 × 10 $\mu s$ |
| MD in M2TM-AH POPC:POPS | 2L0J | 16 | 4768 | 4:1 | 2 × 10 $\mu s$ |
| MD in M2TM-AH POPC:POPS:cholesterol | 2L0J | 16 | 4752 | 3:1:1 | 2 × 10 $\mu s$ |
| MD in M2TM-AH plasma mimetic membrane. Upper leaflet POPC:DOPC:POPE:DOPE: DPSM:DPG3:CHOL Lower leaflet POPC:DOPC:POPE:DOPE:POPS:DOPS :POP2:CHOL | 2L0J | 16 | 4852 | Upper 20:20:5:5: 15:10:25 Lower 5:5:20:20:8:7: 10:25 | 2 × 10 $\mu s$ |
| MD in M2TM-AH POPC:cholesterol | 2L0J | 3 | 546 | 4:1 | 2 × 50 $\mu s$ |
| MD in M2TM-AH POPC:POPS:cholesterol | 2L0J | 3 | 617 | 3:1:1 | 3 × 10 $\mu s$ |
| PMF(US) in M2TM-AH POPC | 2L0J | 2 | 546 | 1 | 518.4 $\mu s$ |
| PMF(US) in M2TM-AH POPC:cholesterol | 2L0J | 2 | 546 | 4:1 | 518.4 $\mu s$ |

| Simulation atomistic | Number of lipids | Bilayer composition | Repeats × duration |
|---|---|---|---|
| MD in dimer of M2TM-AH channels | 546 | 4:1 | 3 × 500 ns |
| MD in dimer of M2TM-AH channels with three cholesterols in between M2 | 546 | 4:1 | 3 × 500 ns |

[a] see Methods section

**Supplementary Table 2.** Cholesterol kinetic parameters for the binding sites 1 (or 4), 2 (or 3), 5 (or 6) [a] defined in Fig. 7, calculated using PyLipID [71] library (dual cutoffs 4.75-7.5 Å) from 2 × 50 $\mu s$ CG MD simulations with Martini force field. [64,127,128,135]

| | Top Leaflet | | Bottom Leaflet |
|---|---|---|---|
| | Site 1 or 4 | Site 2 or 3 | Site 5 or 6 |
| Binding site residence time | 4.669 $\mu$s | 9.802 $\mu$s | 15.410 $\mu$s |
| Site occupancy | 66.736 % | 92.050% | 88.548 % |
| Site surface area ($nm^2$) | 7.733 | 17.122 | 13.155 |
| $k_{off}$ | 0.214 $\mu s^{-1}$ | 0.102 $\mu s^{-1}$ | 0.065 $\mu s^{-1}$ |
| $R^2$ | 0.9917 | 0.9897 | 0.9765 |
| Average # cholesterols in site | 1.348 | 1.988 | 2.109 |

[a] see Methods section

**Supplementary Table 3.** Cholesterol kinetic parameters for the binding sites 1 (or 4), 3, 5 (or 6) defined in Fig. 5 including 20% POPS in each leaflet. Sites were calculated using PyLipID [71,] library (dual cutoffs 4.75-7.5 Å) from 3 × 10 μs CG MD simulations with Martini force field. Binding site residence times are capped to 10 μs.

| Cholesterol sites incl. 20% POPS | Top Leaflet | | Bottom Leaflet |
|---|---|---|---|
| | Site 1 or 4 | Site 3 | Site 5 or 6 |
| Binding site residence time | 3.034 $\mu$s | 1.779 $\mu$s | 8.662 $\mu$s |
| Site occupancy | 96.124 % | 59.681 % | 92.667 % |
| $k_{off}$ | 0.330 $\mu s^{-1}$ | 0.562 $\mu s^{-1}$ | 0.067 $\mu s^{-1}$ |
| $R^2$ | 0.9520 | 0.9941 | 0.9851 |
| Average # cholesterols in site | 2.132 | 1.145 | 1.507 |

**Supplementary Table 4.** Descriptive statistics gaussian curvature values shown in Fig 9 and Fig S6. Data is averaged over the final 2 μs and over two independent repeat simulations.

| | Protein-free | | M2TM | | | M2TM-AH | | | |
|---|---|---|---|---|---|---|---|---|---|
| Gaussian curvature (Å$^{-2}$) | POPC | POPC/Chol | DMPC | POPC | POPC/Chol | DMPC | POPC | POPC/Chol | POPC/POPS/Chol |
| Minimum | -0.221185 | -0.569526 | -0.272171 | -0.253168 | -0.244917 | -0.240787 | -0.301885 | -1.35670 | -0.321659 |
| Maximum | 0.0345786 | 0.588636 | 0.295888 | 0.0341831 | 0.0462298 | 0.210733 | 0.577205 | 0.951558 | 1.04557 |
| Range | 0.255763 | 1.15816 | 0.568059 | 0.287351 | 0.291147 | 0.451520 | 0.879091 | 2.30825 | 1.36723 |
| Mean | -0.0230189 | -0.0320807 | -0.0291107 | -0.0265420 | -0.0271631 | -0.0265916 | -0.0274372 | -0.0366976 | -0.0309849 |
| Std. Deviation | 0.0486328 | 0.0866705 | 0.0635267 | 0.0507486 | 0.0517711 | 0.0542209 | 0.0784003 | 0.142040 | 0.0954490 |

### Supplementary Figures

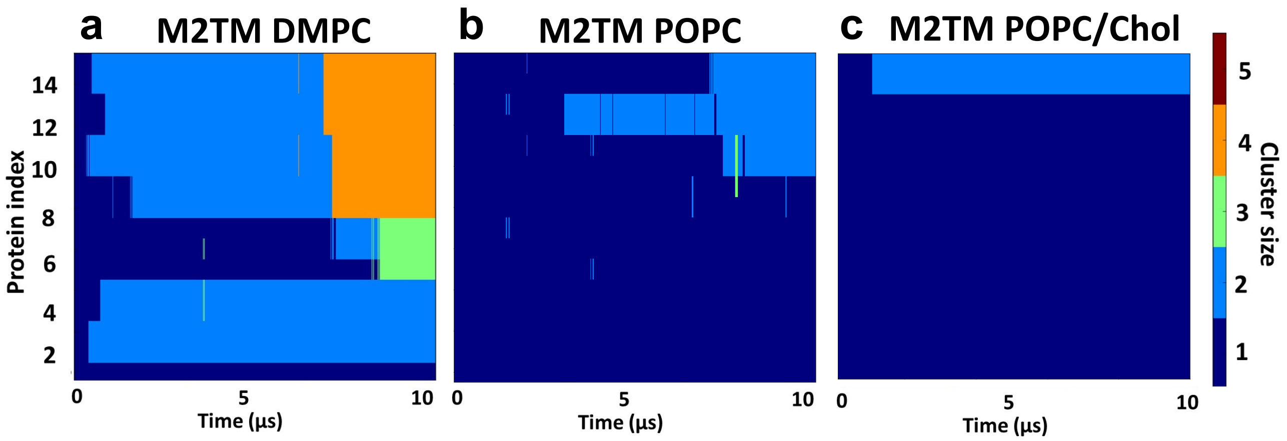

**Supplementary Fig. 1.** The formation of clusters of M2 channels over the duration of 10 *μs*-CG MD simulations with Martini force field [64,127,128,135] is illustrated for simulations of 16 copies of the M2 channel protein in bilayers: (a) M2TM in DMPC, (b) M2TM in POPC, and (c) M2TM in POPC/Chol. The cluster size is indicated by colour (blue for single M2 channels, cyan for dimers of M2 channels, green for trimers, orange for tetramers, and brown for pentamers).

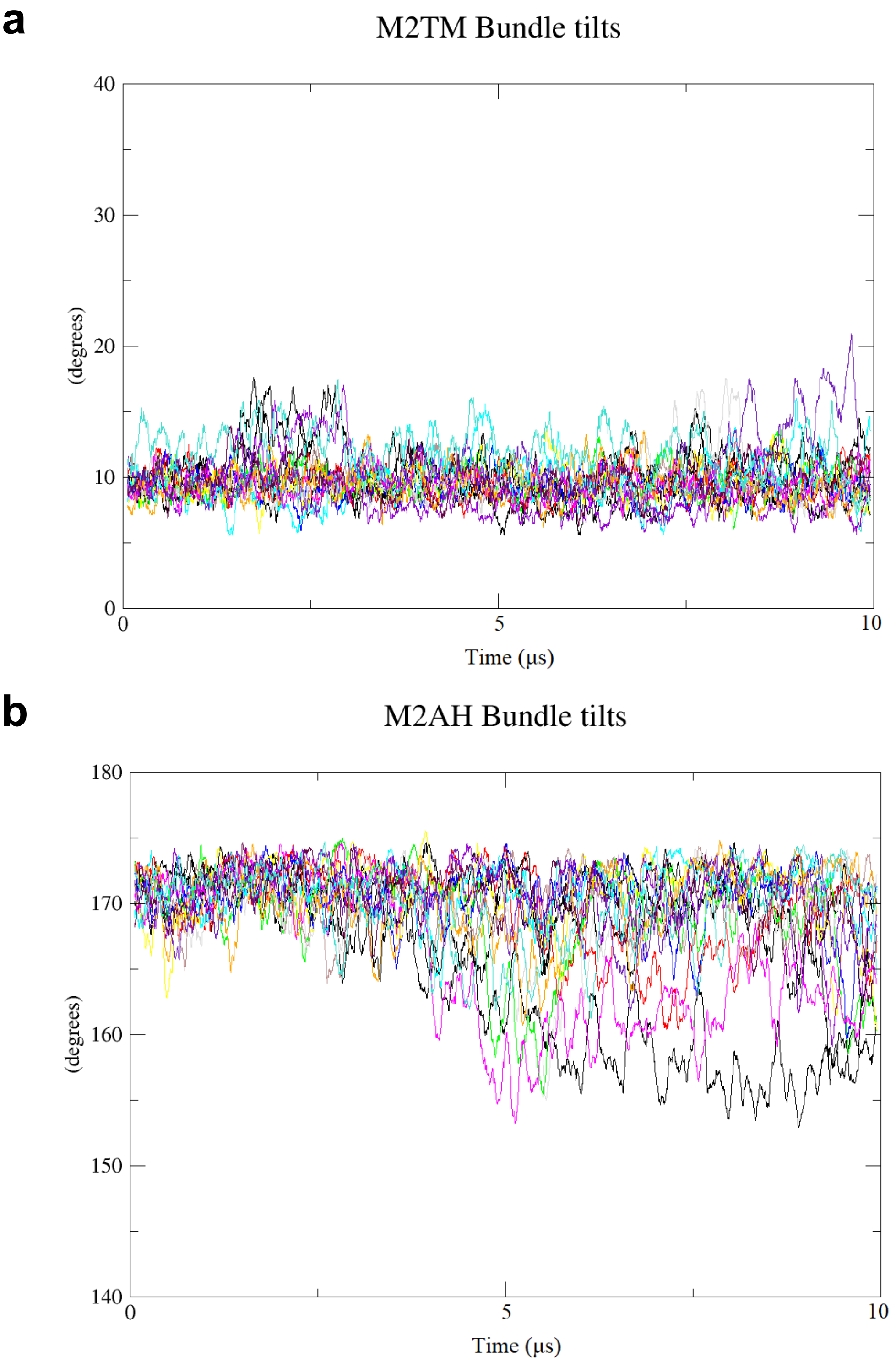

**Supplementary Fig. 2.** Evolution of the angle between M2TM or M2TM-AH bundle axis and the membrane normal over the simulation time from 10 $\mu s$-CG MD simulations with Martini force field [64,127,128,135] of 16 copies of the M2 channel protein in bilayers. Each colour corresponds to one M2TM or M2TM-AH protomer of the systems DMPC or POPC, respectively.

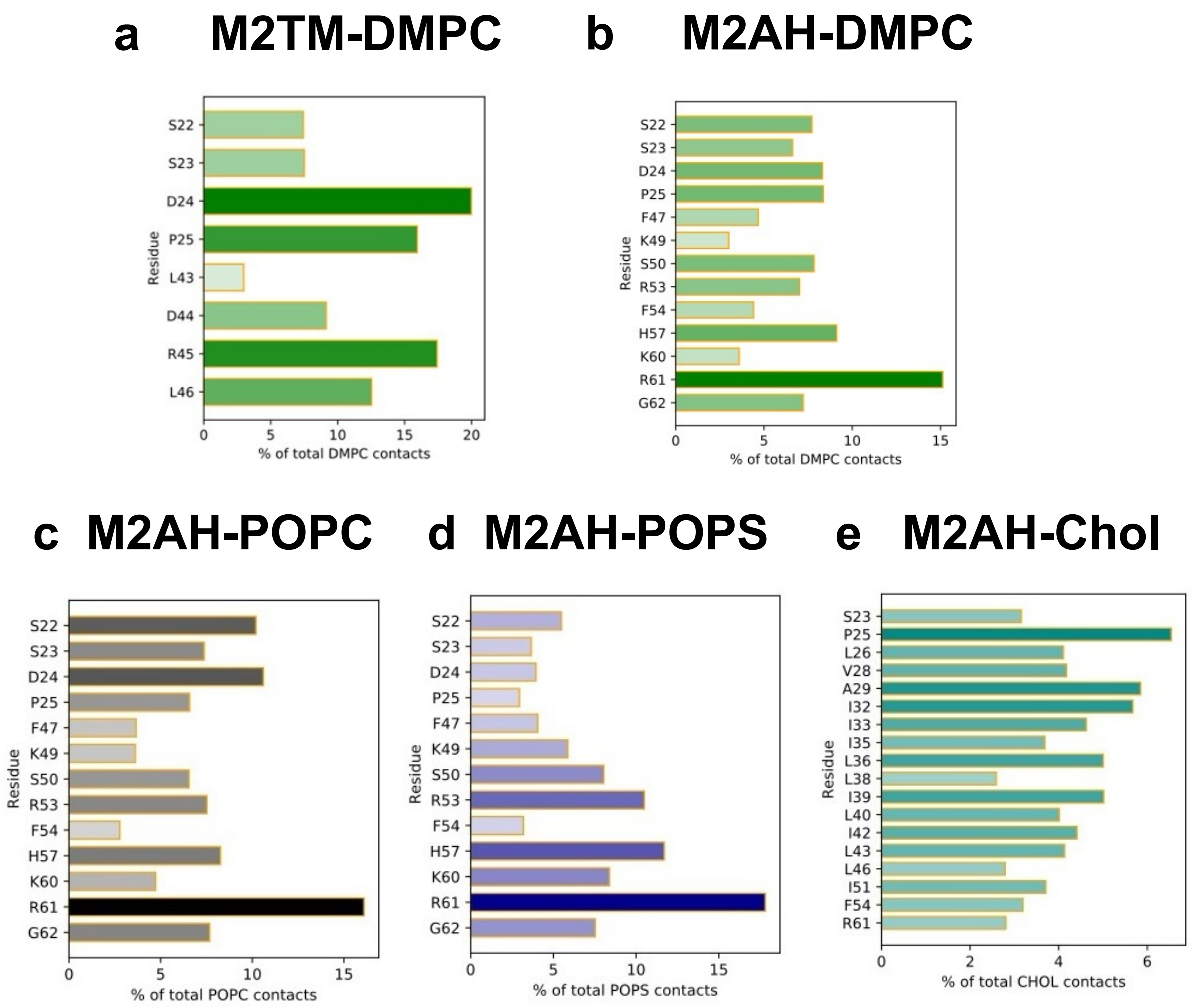

**Supplementary Fig. 3.** Protein-lipid interaction plots associated with Fig. 4. The bars are coloured according to the frequency of residue interactions with phospholipid headgroups (PO4 beads) or with cholesterol. Heatmaps show lipid contacts in simulations of **a**. M2TM with DMPC lipid headgroups; **b**. M2TM-AH with DMPC lipid headgroups; **c**. M2TM-AH with POPC lipid headgroups; **d**. M2TM-AH with POPS lipid headgroups and **e**; M2TM-AH with cholesterol.

- Cholesterol system

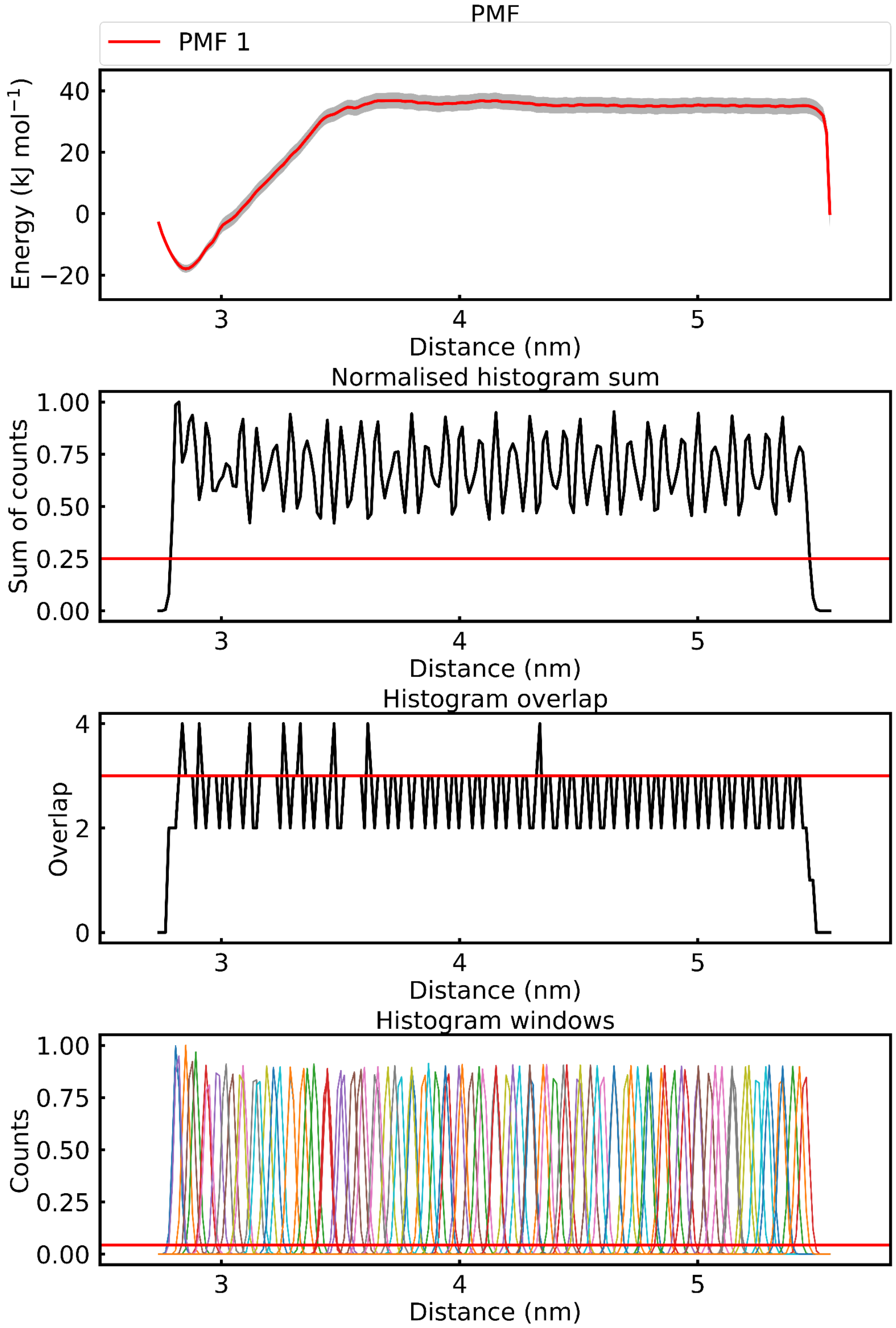

**Supplementary Fig. 4.** PMF calculation for the dissociation of dimer of M2TM-AH channels in POPC (referring to Fig. 8 -cholesterol PMF). PMF plots and histograms derived with WHAM [146,147] analysis after umbrella sampling along the CV (protomer center of mass distance) with the Martini force field [64,127,128,135] . From bottom to top are shown the histograms of US windows, neighbouring histogram overlap needed for sampling, and the normalized histograms.

+ Cholesterol system

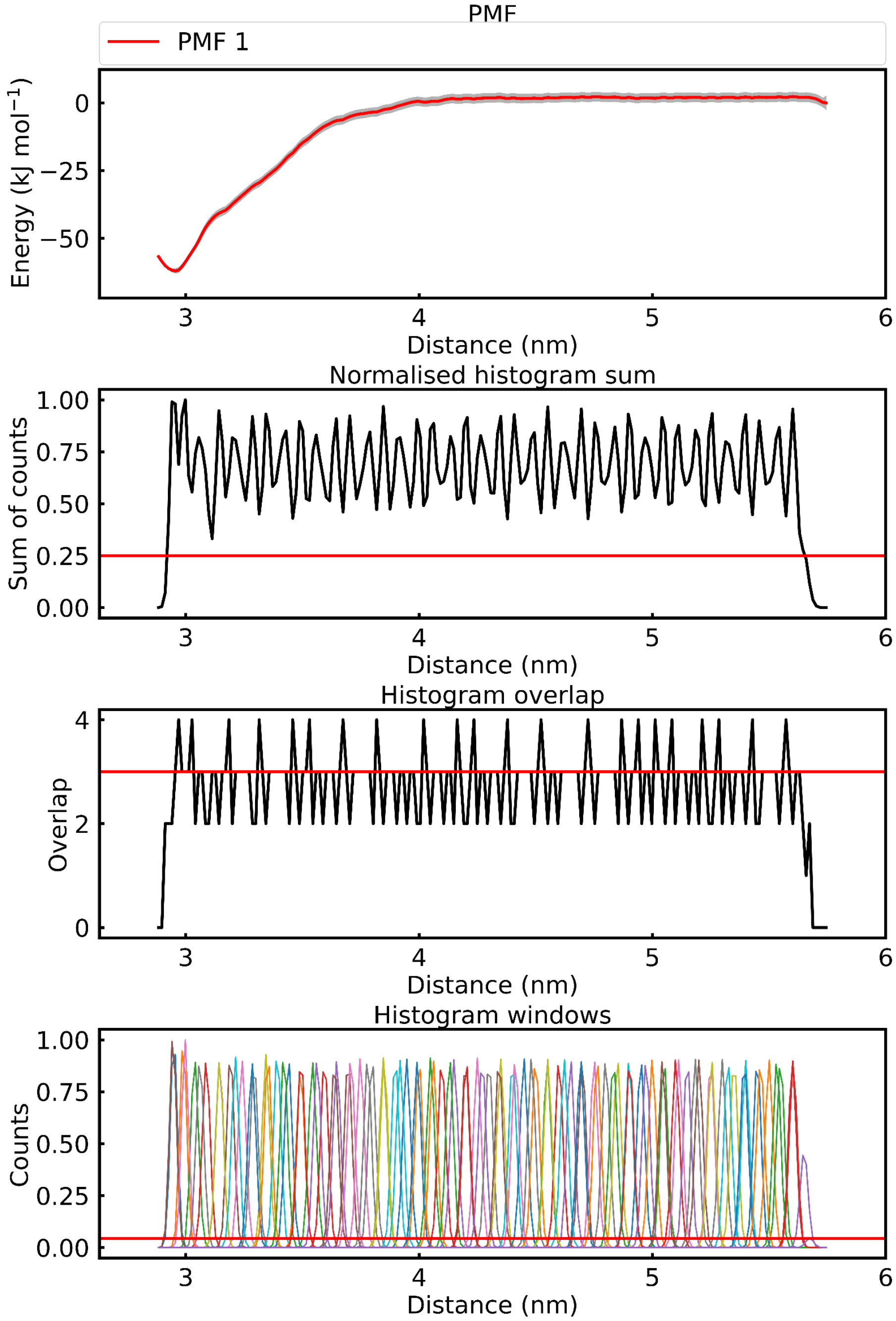

**Supplementary Fig. 5.** PMF calculation for the dissociation of dimer of M2TM-AH channels in POPC/cholesterol with three cholesterols occupying the interfacial binding sites between M2TM-AH protomers in the dimer discussed in the main text (referring to Fig. 8 +cholesterol PMF). PMF plots and histograms derived with WHAM [146,147] analysis after umbrella sampling along the CV (protomer center of mass distance) with the Martini force field. [64,127,128,135] From bottom to top are shown the histograms of US windows, neighbouring histogram overlap needed for sampling, and the normalized histograms.

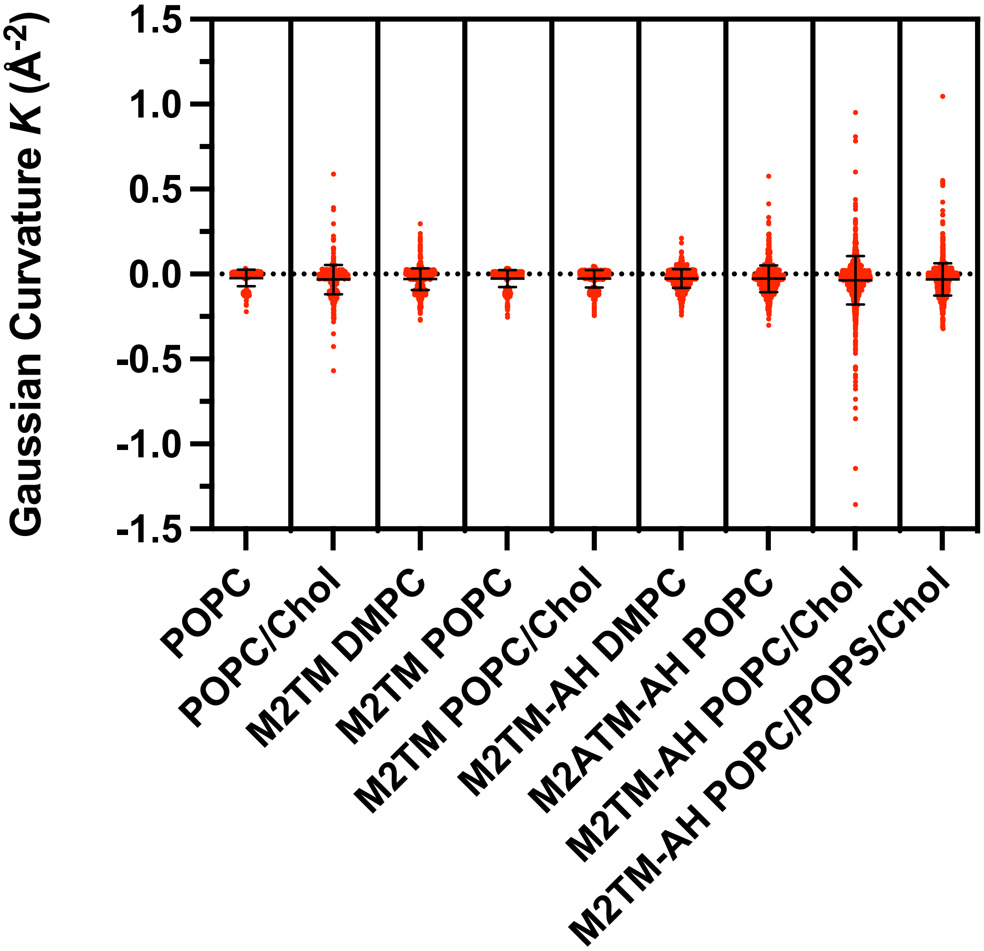

**Supplementary Fig. 6.** Scatter plot of gaussian curvature values shown in Fig 9. Data points are shown as red dots and mean and standard deviation with black bars.

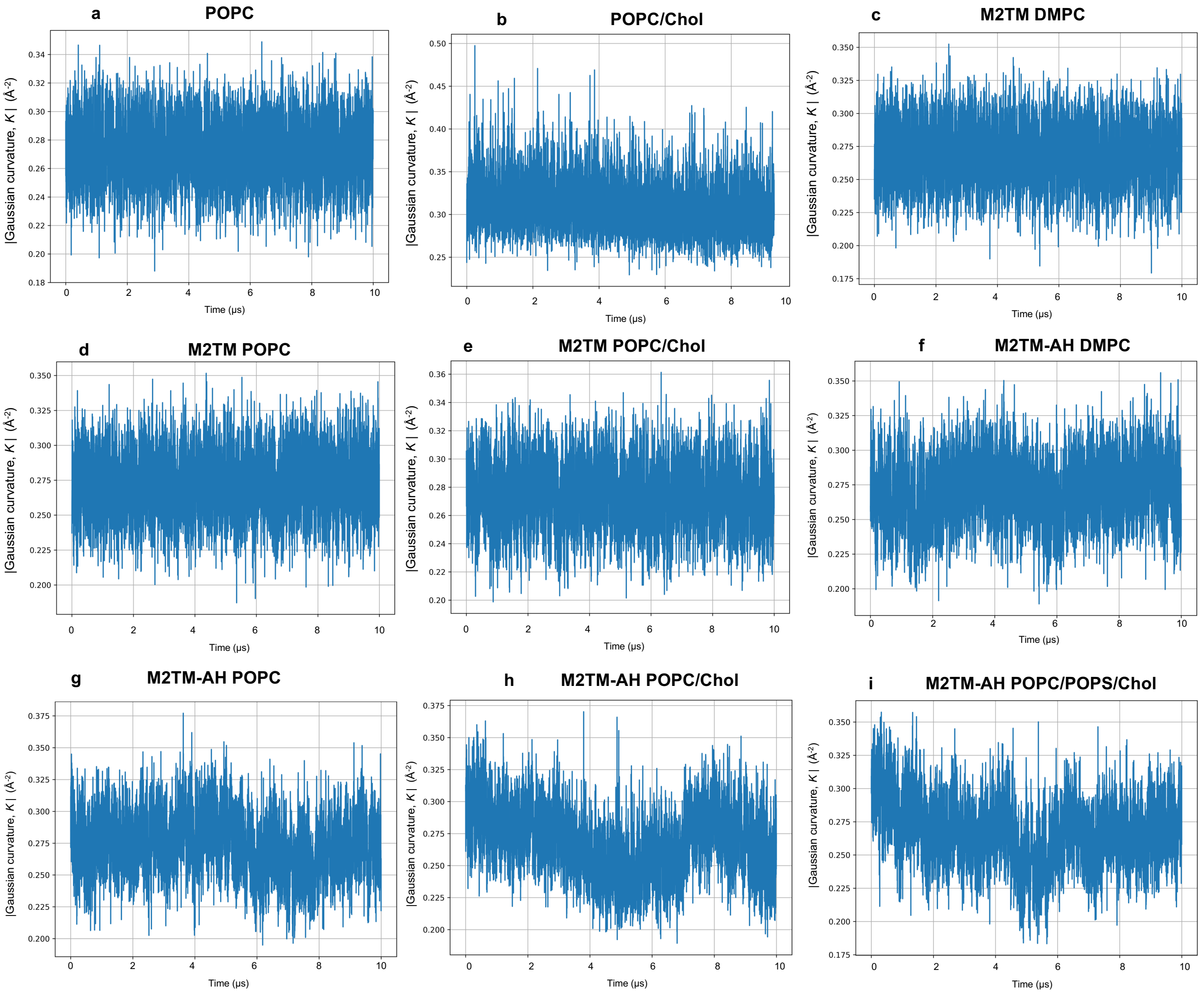

**Supplementary Fig. 7.** Absolute values of the mean Gaussian curvature (|$K$|) from different systems (protein-free, M2TM, M2TM-AH) as a function of trajectory time (μs) from CG MD simulation with the Martini force field. [64,127,128,135] Each data point represents the average (over discretized grids, *i,j*) per frame of the absolute Gaussian curvature. Absolute values were tracked per frame to avoid cancelling out of the curvature across a bilayer, which has positive and negative local values. Thus, the higher the absolute value, the higher degree of total Gaussian curvature a bilayer has. Systems: **a**. POPC; **b**. POPC/cholesterol; **c.** M2TM/DMPC; **d.** M2TM/POPC; **e.** M2TM/POPC/cholesterol; **f.** M2TM-AH/DMPC; **g.** M2TM-AH/POPC; **h.** M2TM-AH/POPC/cholesterol; **i.** M2TM-AH/POPC/POPS/cholesterol.

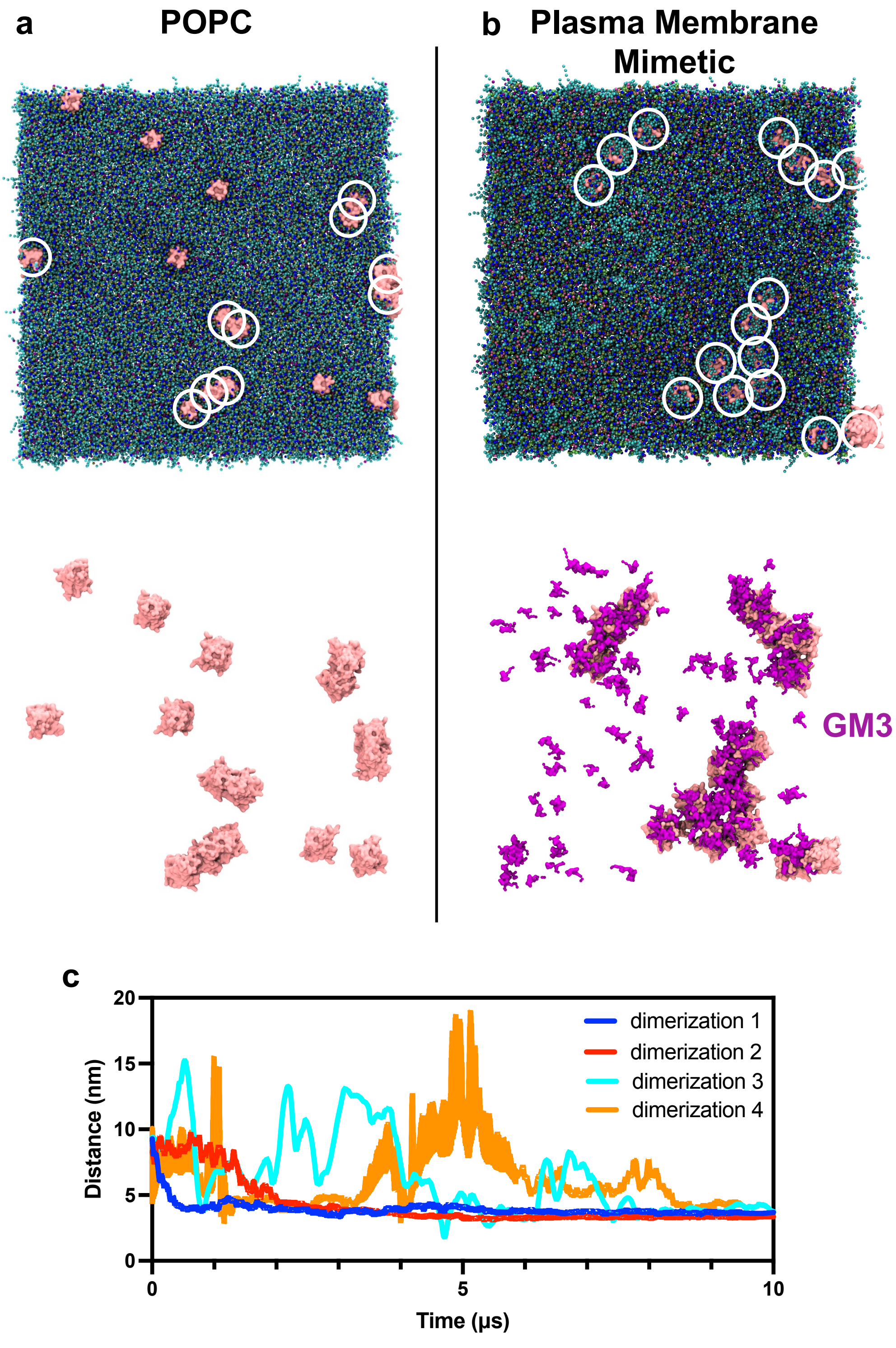

**Supplementary Fig. 8. a**. Top views of the last frame from CG MD simulations with Martini force field [64,127,128,135] of the M2TM-AH POPC system (top figure; protein backbone in pink and membrane in blue and green colours, bottom figure; same as top but without membrane for the sake of clarity). M2TM-AH protomers that form dimers are shown in white circles. **b**. Top views of the last frame of the M2TM-AH plasma mimetic membrane system (top figure; protein backbone in pink and membrane in blue and green colours, bottom figure; same as top but without membrane for the sake of clarity, GM3 lipids gathered around M2TM-AH protomers, are depicted in purple). M2TM-AH protomers that form oligomers are shown in white circles, with a white line connecting them **c**.

Distance measurement in the M2TM-AH plasma mimetic membrane system; distance (nm) between the COM of protomers in a dimer of M2TM-AH channels as a function of time (μs). Despite the GM3 shells resulting in a higher COM distance between protomers, the analysis shows that clusters are stable once formed.

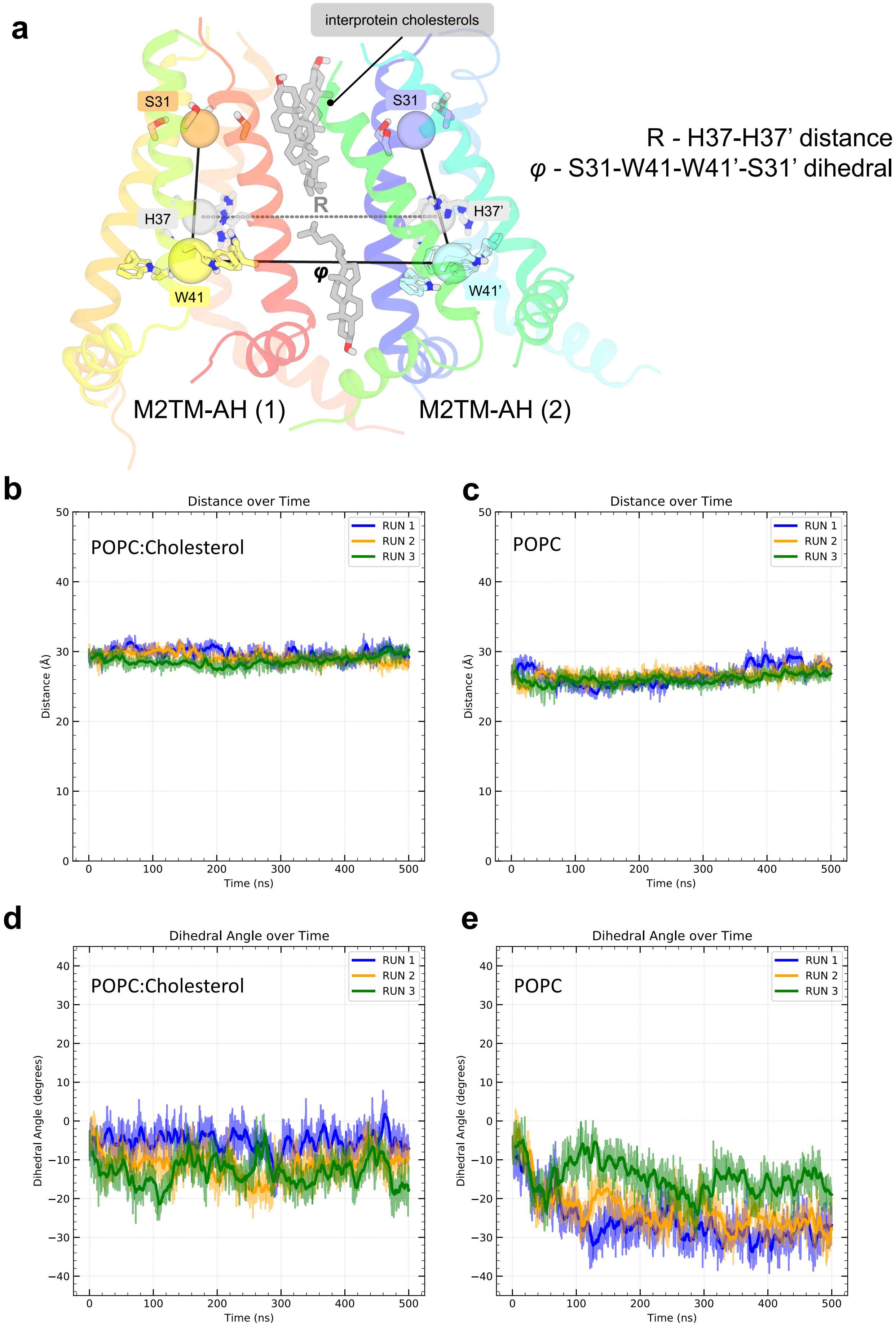
a
interprotein cholesterols
S31
S31
R - H37-H37' distance
φ - S31-W41-W41'-S31' dihedral
R
H37
H37'
φ
W41
W41'
M2TM-AH (1)
M2TM-AH (2)
b
Distance over Time
POPC:Cholesterol
RUN 1
RUN 2
RUN 3
Distance (Å)
Time (ns)
c
Distance over Time
POPC
Distance (Å)
Time (ns)
d
Dihedral Angle over Time
POPC:Cholesterol
Dihedral Angle (degrees)
Time (ns)
e
Dihedral Angle over Time
POPC
Dihedral Angle (degrees)
Time (ns)

**Supplementary Fig. 9.** Stability analysis of M2TM-AH dimers in POPC and POPC:Cholesterol (4:1) from AA MD simulations with the CHARMM36m force field.[79] **a**. Structure-based definitions of tetrad of COMs used for interprotomer distance and relative rotation tracking [H37 residues from protomer (1) and H37′ from (2) defined two COMs and the COM-COM distance (R) was calculated; The S31 and W41 tetrads from protomer (1) and W41′ and S31′ tetrads from protomer (2) defined 4 points for the dihedral, $\varphi$, tracking]; M2TM-AH dimer **b.** distance R over time in POPC/Chol; **c.** distance R over time in POPC; **d.** dihedral $\varphi$ over time in POPC/Chol; **e.** dihedral $\varphi$ over time in POPC. Three independent repeats were run for each system (run 1: blue, run2: orange, run 3: green). The trajectory data are plotted per frame (100 ps/frame) in transparent lines and the rolling average of each metric is plotted in opaque. Rolling averages were calculated using a window size of 100 frames.

Figures

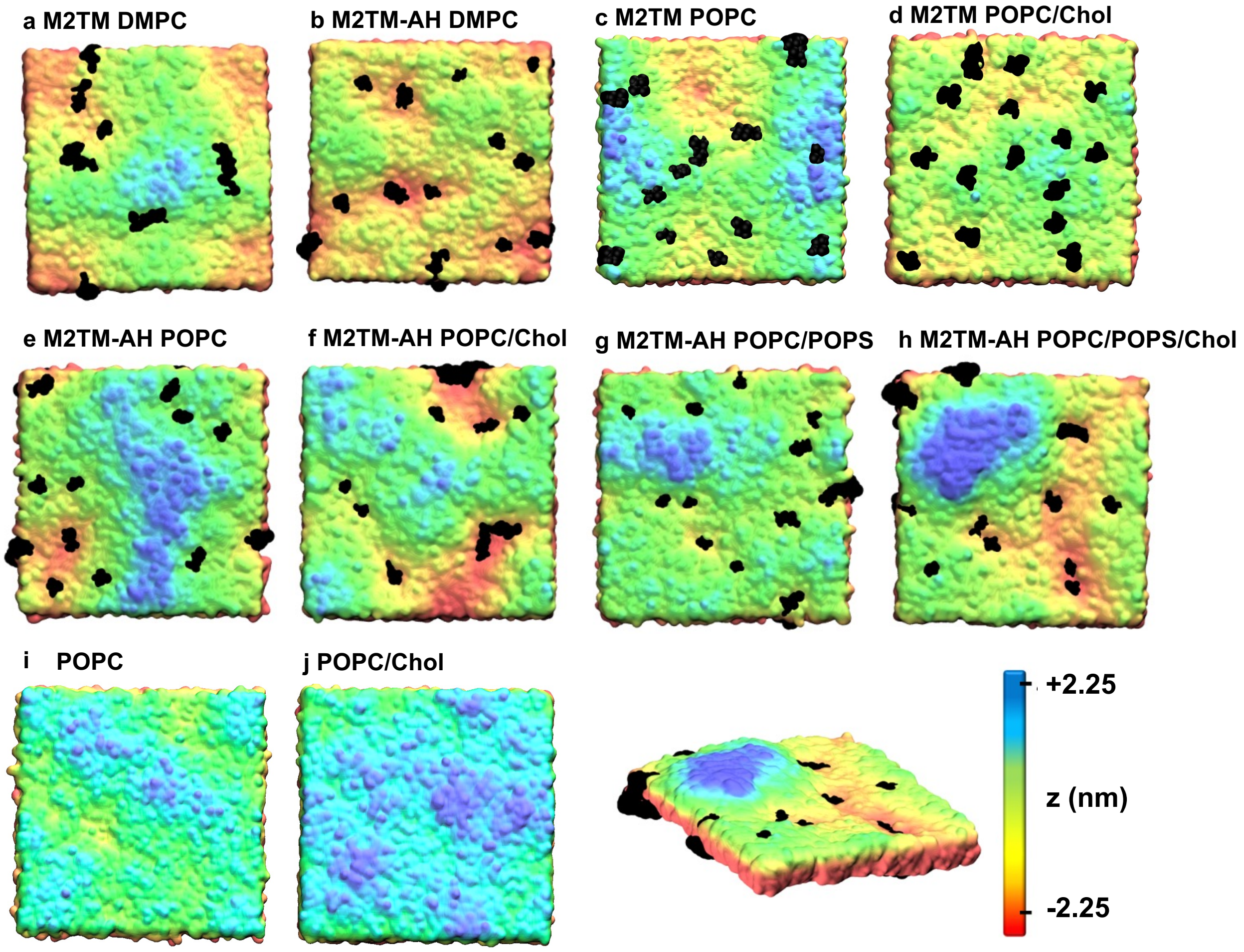

**Supplementary Fig. 10.** Snapshots of local membrane deformation and formation of M2 channel clusters from CG MD simulation with Martini force field. [64,127,128,135] Representative simulation snapshots (taken at $t = 10\ \mu s$) of 16-copy M2 channels are shown (top view) from CG MD simulations of **a**. M2TM in DMPC; **b**. M2TM-AH in DMPC; **c**. M2TM in POPC; **d**. M2TM in POPC/cholesterol; **e**. M2TM-AH in POPC; **f**. M2TM-AH in POPC/cholesterol; **g**. M2TM-AH in POPC/POPS; **h**.

M2TM-AH in POPC/POPS/Chol; **i**, **j**. Pure POPC, POPC/cholesterol bilayers, respectively. Non-protein surfaces are presented as colour heatmaps according to local z value (midplane is taken as reference z = 0; red is assigned to z < 0 and to blue for z > 0, ranging from z = -2.25 nm to z = +2.25 nm). Protein surfaces are in black for clarity. There is visual correlation between M2TM-AH clusters and negative mean curvature (*H*, i.e. membrane "valleys"). The term was used to avoid confusion from discussing two curvature types (mean, *H*, and Gaussian, *K*).

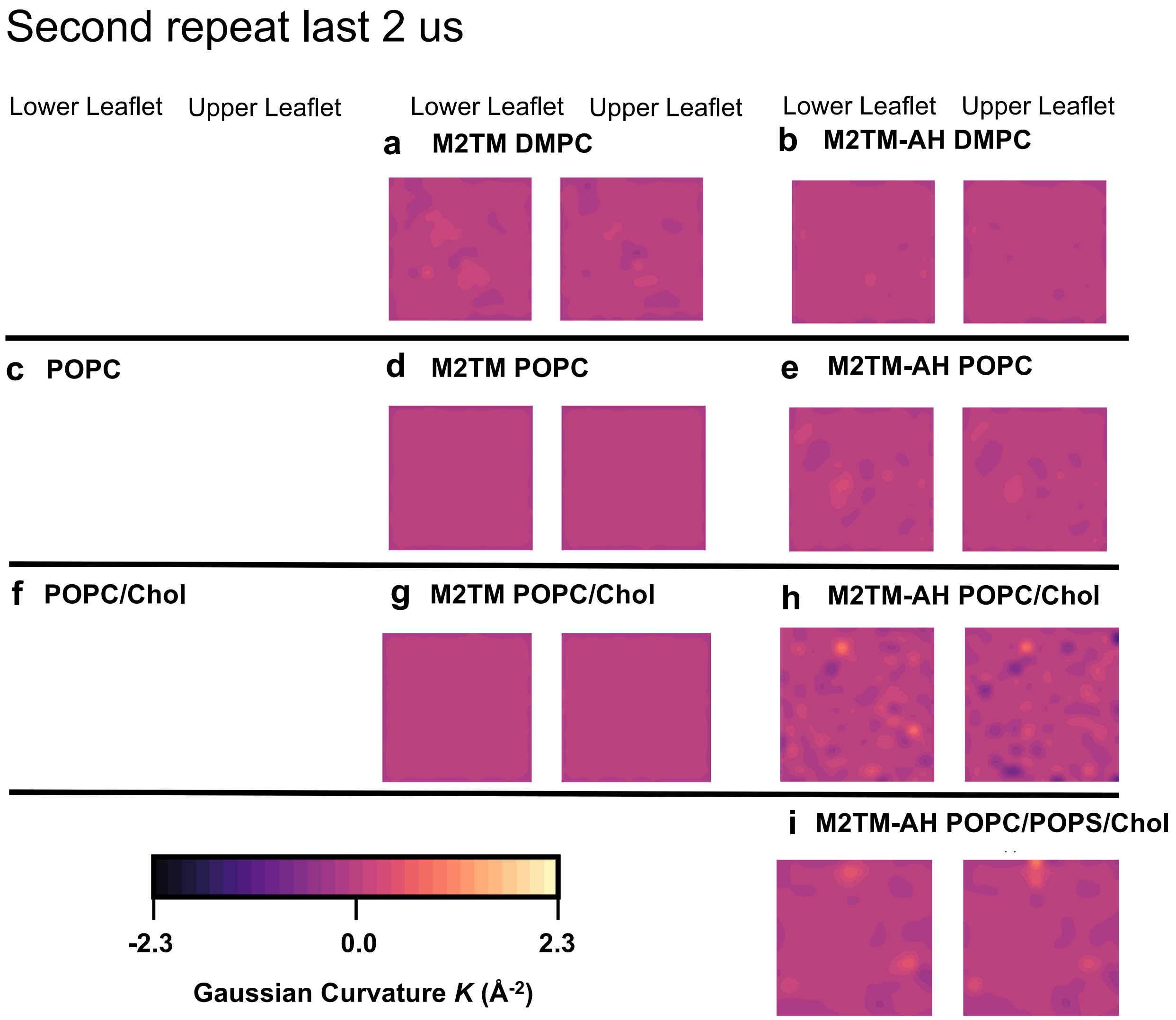

**Supplementary Fig. 11.** Gaussian curvature (*K*) induced by M2 constructs (±AHs) and cholesterol from 10 $\mu$s CG MD of 16 copies of the M2 constructs and protein-free control bilayers with the Martini force field. [64,127,128,135] The *K* heatmaps are averaged over the final 2 $\mu$s from a single replicate. Results for the first replicate are shown in main text Fig. 9.

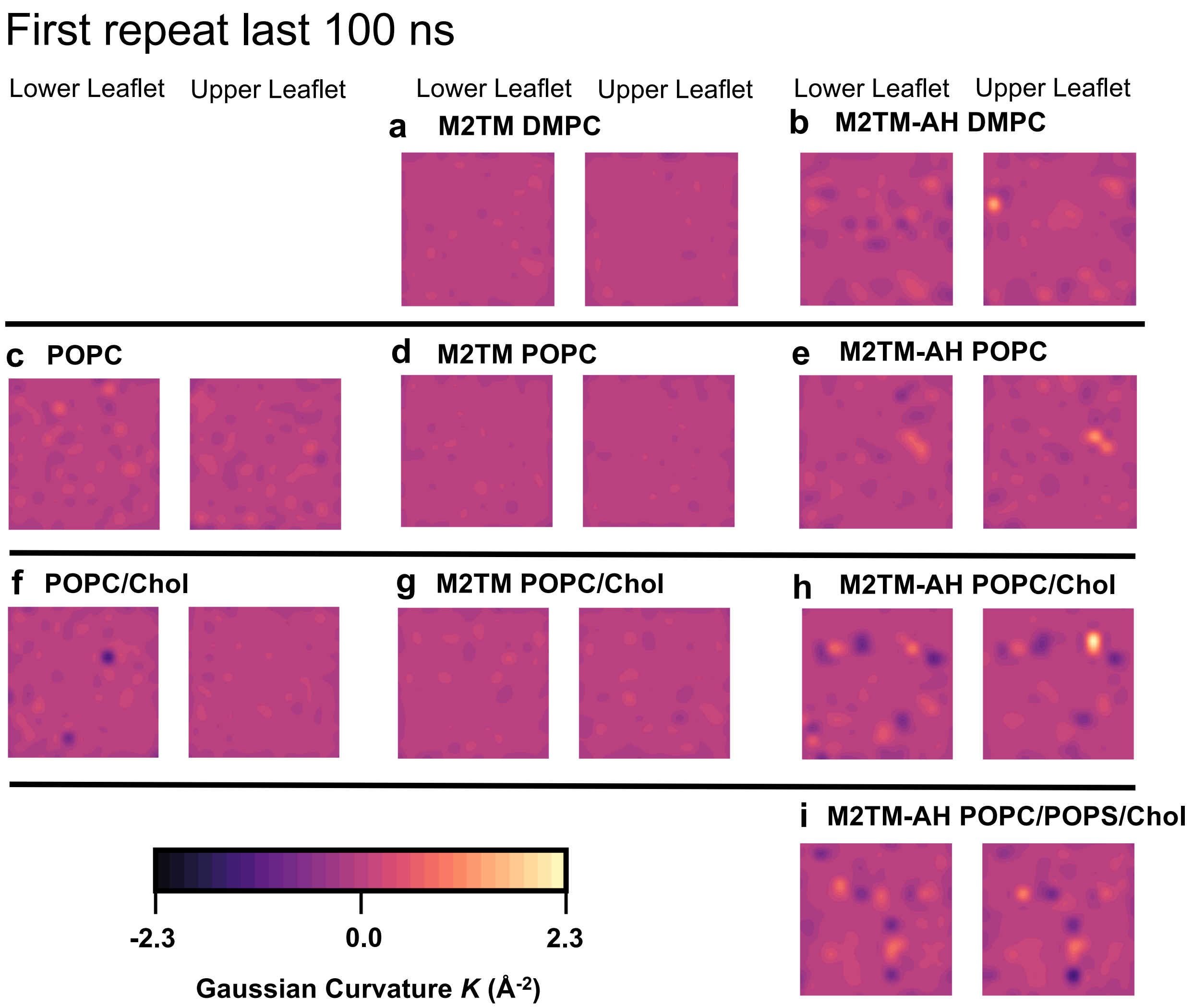

**Supplementary Fig. 12.** Gaussian curvature ($K$) induced by M2 constructs (±AHs) and cholesterol from 10 $\mu$s CG MD of 16 copies of the M2 constructs and protein-free control bilayers with the Martini force field. [64,127,128,135] The $K$ heatmaps are averaged over the final 100 ns to accentuate the temporally resolved membrane undulations. Data extracted from the first replicate simulations.

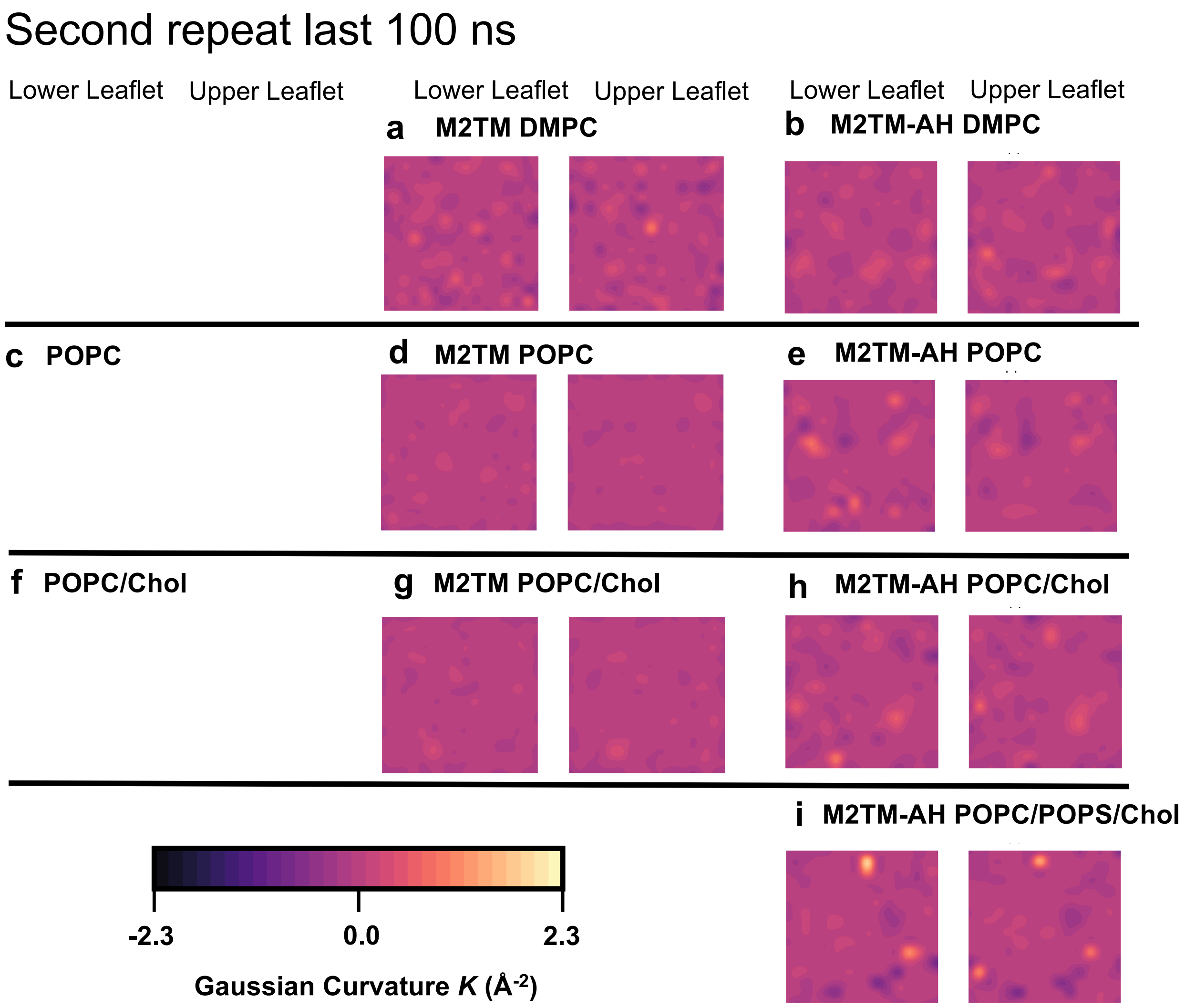

**Supplementary Fig. 13.** Gaussian curvature (*K*) induced by M2 constructs (±AHs) and cholesterol from 10 $\mu$s CG MD of 16 copies of the M2 constructs and protein-free control bilayers with the Martini force field. [64,127,128,135] The *K* heatmaps are averaged over the final 100 ns to accentuate the temporally resolved membrane undulations. Data extracted from the second replicate simulations.
